# Structural probing of RNA hairpins quantifies protein occupancy on RNA and links it to function in human cells

**DOI:** 10.64898/2026.01.06.698062

**Authors:** Abby R. Thurm, Tanya H. Nguyen, Jesus Damian Torres Campos, Matthew Mansfield, Lacramioara Bintu, Lauren D. Hagler

## Abstract

RNA structures that form inside living cells influence processes ranging from translation to RNA decay, many of which are controlled by RNA-binding proteins (RBPs). Because RBP specificity depends on both local RNA structure and sequence motifs, traditional pulldown-based methods often obscure the structural context of bound RNAs. Here, a quantitative framework based on dimethyl sulfate mutational profiling and sequencing (DMS-MaPseq) is introduced that jointly measures RNA structure and protein binding at single-nucleotide resolution in human cells, enabling estimation of effective RNA-protein affinities and fractional occupancy directly in cells. Application of this approach to a 1,600-member library of MS2 hairpin mutants reveals that RNA folding is the strongest determinant of MS2 coat protein (MCP) recognition, and that stable MS2 structures must form for MCP to bind its target sequence. MCP also shows strong preference for its consensus loop sequence while displaying minimal dependence on stem length beyond ten base pairs or on stem GC content. Incorporation of an inducible degron fused to MCP allows precise tuning of intracellular protein concentrations analogous to those of many endogenous RBPs and show that DMS reactivity changes can be used to infer binding specificities across a subsaturating regime. A quantitative occupancy framework further shows that inferred fraction-bound values accurately predict how efficiently MCP fused to a downregulatory-domain drives RNA degradation. Together, these results establish a generalizable approach for measuring RBP-RNA affinities with structural resolution in living cells, dissecting how sequence and structure contribute to RBP recognition, and quantitatively linking occupancy to functional output.

## INTRODUCTION

RNA is a central regulator of biological function that plays important roles in a wide range of cellular processes.^1,2^ By folding into dynamic secondary and tertiary structures and associating with RNA-binding proteins (RBPs), RNAs dictate complex interactions that control such fundamental processes as translation, splicing, and localization.^3–7^ This network of interactions represents a crucial layer of post-transcriptional control that fine-tunes gene expression in response to external signals.^8,9^ In order to fully understand the extent to which RNAs regulate overall cellular behavior, it is first necessary to determine how much of a specific RNA-RNA or RNA-protein interaction, in terms of fractional stability or occupancy, is required to trigger a measurable biological function. In principle, this would involve measuring the affinity (*K*_d_) and bound fraction of an RNA-RBP complex directly in its native cellular context, where both RNA structure and protein availability are physiologically relevant. However, such measurements remain experimentally challenging with existing genomic approaches, as they require control over RNA and/or protein concentration and direct quantification of the fraction of molecules that are bound.^10,11^

High-throughput methods such as enhanced crosslinking and immunoprecipitation (eCLIP)^12^ have been instrumental in identifying RNA-protein interactions across the transcriptome and in defining consensus binding motifs for RBPs.^13^ By aggregating the sequences of enriched RNA fragments, eCLIP enables construction of position-weight matrices that describe sequence preferences and support inference of relative binding specificity.^14^ However, these bulk measurements provide no information about the extent of binding at individual RNA sites. Because eCLIP signals reflect population-averaged enrichment across thousands of transcripts, they cannot report what fraction of a given RNA is actually bound by an RBP at endogenous levels in cells, nor how RNA structure modulates that occupancy.^15^ As a result, while eCLIP robustly identifies where RBPs can bind, it cannot determine how much binding is required to drive regulatory function at a specific RNA locus.^16,17^ Addressing this gap requires an approach that can quantify RNA-protein interactions at defined sites in living cells while simultaneously reporting on the underlying RNA structural state that enables, or restricts, binding.

RNA chemical probing has been traditionally used to predict the secondary structure of RNAs transcriptome-wide.^18^ Methods like DMS-MaPseq (Dimethyl Sulfate Mutational Profiling with Sequencing)^19^ and SHAPE (Selective 2’-Hydroxyl Acylation analyzed by Primer Extension)^20^ use chemical probes like dimethylsulfate and anhydrides or imidazolides, respectively, to selectively modify RNA based on its structural state. During DMS-MaPseq, dimethylsulfate is used to methylate the Watson-Crick-Franklin face of adenine (N1) and cytosine (N3) nucleotides when they are unpaired and solvent-exposed, enabling discrimination between structured and unstructured regions of RNA.^21^ Methylations are converted to mutations during reverse transcription and detected by next-generation sequencing.^22,23^ Mutational profiles can be used to constrain secondary structure prediction and discover novel RNA structures, often with functional consequences.^18^ Despite its power, RNA chemical probing has largely been applied to determine where RNA is folded, rather than how much of an RNA population occupies a given structural state or how that stability relates to downstream function.^24^ Critically, RNA folding and protein binding are tightly coupled processes,^25,26^ yet existing applications of DMS-MaPseq do not explicitly link structural ensembles to RBP occupancy. This limitation motivates the development of approaches that leverage chemical probing not only to report RNA structure, but also to quantitatively measure RNA-protein interactions in their native cellular context.

The differential reactivity of dimethyl sulfate based on the local environment of the RNA is not limited to constraining models for structure prediction; it can also be used to map binding events of other RNAs, proteins, and small molecule ligands by detecting changes in RNA accessibility due to direct shielding or a change in conformation.^21,27–33^ Building upon this principle, we developed a quantitative framework for measuring structural stability and occupancy simultaneously in human cells using RNA chemical probing. A systematic library in which each position of an RNA is individually mutated enabled direct assessment of how each single-nucleotide change influences local folding, protein binding, and downstream functional outcomes. The MS2 hairpin-MS2 coat protein (MCP) system was selected as a model system because it has a well-defined motif and is well-characterized structurally such that the MCP-interacting bases in the MS2 hairpin are known (**Figure 1A**).^34,35^ DMS-MaPseq can effectively and sensitively measure MCP binding to MS2 hairpins in cells, revealing that stable hairpin formation is a prerequisite for MCP binding and that single-nucleotide changes in the loop can quantitatively modulate affinity. Incorporation of a degron-controlled MCP variant further enabled modulation of intracellular protein levels to values similar to those of endogenous RBPs, allowing characterization of binding preferences under subsaturating conditions. Finally, a quantitative framework for MCP occupancy in living cells showed that these predictions correlate with the functional output of a downregulatory RBP, establishing a formative link between occupancy and function through a base-resolved method for measuring RBP binding in cells.

**Figure 1.**
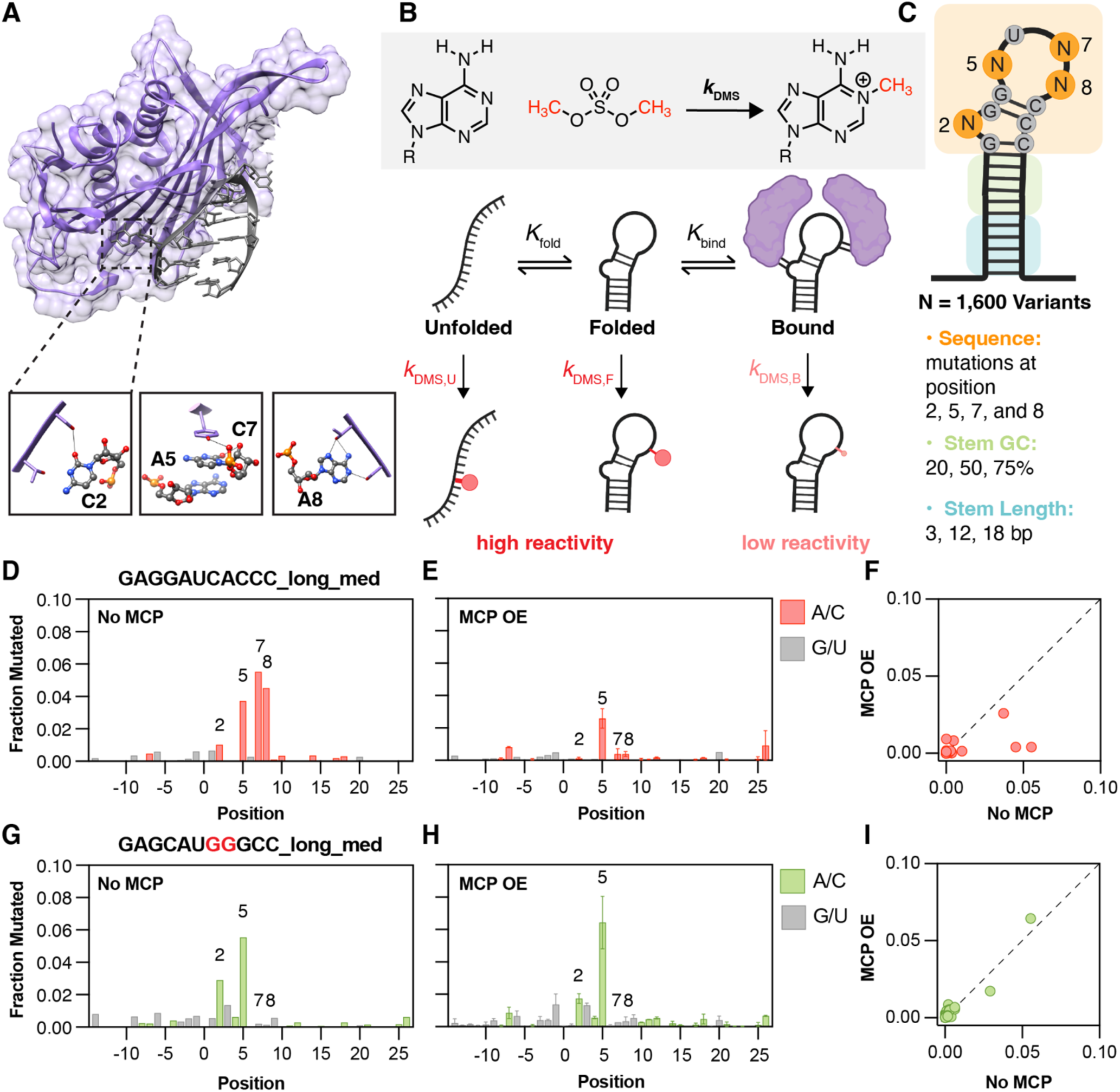
DMS-MaPseq measurements of RNA-protein binding. (**A**) Top, X-ray structure of MS2 coat protein (MCP, orange) recognizing and binding to the canonical MS2 stem-loop (grey, PDB: 2IZM).^39^ Bottom, specific MCP residues make direct contact with designated places on the MS2 hairpin. Tyrosine 85 touches the first and third positions of the tetraloop (5 and 7 by our numbering scheme), serine 47 touches the bulge nucleotide at position 2, and threonine 45/serine 47 make contact with the fourth position of the tetraloop (A8). (**B**) Thermodynamic scheme for RNA folding and binding. Kinetic scheme for modification by dimethyl sulfate (DMS), which methylates the N1 position of adenine or the N3 position of cytosine bases. We assume that RNA can exist in three states: unfolded, folded, and folded+bound (bound). The rate of DMS modification at accessible (unpaired) bases is relatively high, leading to high DMS signal under those conditions. When the RNA is completely bound by MCP, the rate of DMS modification at unpaired but bound positions is very low or effectively 0. (**C**) A 1,600 member MS2 stem-loop library varying base identity at positions 2, 7, and 8, stem length, and stem GC content was cloned into the 3’UTR of an mCherry-expressing base plasmid. (**D-E**) DMS reactivity (fraction mutated) at all positions of a wild-type MS2 stem-loop **GAGGAUCACCC** in the absence (**D**) and presence (**E**) of MCP. A and C bases are represented by red bars. G and U bases are represented by grey bars. Errors bars represent the range of two independent replicates. (**F**) Comparison of DMS-MaPseq reactivity at the same positions in the absence (x-axis) and presence (y-axis) of MCP, for the same wild-type stem loop in (**D-E**). Each red circle represents one base averaged over two independent replicates. The line of identity (y=x) is represented by a dashed line. (**G-H**) DMS reactivity (fraction mutated) at all positions of a mutated MS2 stem-loop **GAGCAU<u>GG</u>GCC** in the absence (**G**) and presence (**H**) of MCP. A and C bases are represented by green bars. G and U bases are represented by grey bars. Errors bars represent the range of two independent replicates. (**I**) Comparison of DMS-MaPseq signal at the same positions in the absence (x-axis) and presence (y-axis) of MCP, for the same wild-type stem loop in (**G-H**). Each green circle represents one base averaged over two independent replicates. The line of identity (y=x) is represented by a dashed line.

## RESULTS AND DISCUSSION

### A Systematic MS2 Hairpin Library for Quantitative Analysis of MCP Binding in Human Cells

To establish a method for systematically quantifying RNA–protein binding with single-nucleotide sensitivity while simultaneously measuring RNA structure, a systematic library was generated utilizing the well-studied MS2 hairpin RNA sequence and its cognate protein binder MCP, which are derived from the MCP bacteriophage and are orthogonal to endogenous RNA-protein interactions in human cells.^36,37^ A library of 1,600 MS2 hairpin mutants was designed based on the wild-type consensus sequence **GAGGAU(C/U)ACCC** to dissect the sensitivity of using DMS to detect changes in MCP binding probabilities (**Table S1**). The RNA sequence was systematically varied to modulate each of the contributing thermodynamic and kinetic parameters (*K_f_*_old_: probability of RNA folding, *K*_bind_: probability of protein binding, *k*_DMS_: rate of DMS modification) based on the stability of the resulting RNA structure and MCP binding affinity (**Figure 1B**). Initially, a wild-type MS2 stem-loop^38^ (**GAGGAUCACCC**) was inserted in a variable stem region in the 3’UTR of a coding region for the fluorescent protein mCherry (**Figure 1C**). To modulate *K*_bind_, all possible base mutants of the bulge A (A2), the C in the third position of the tetraloop (C7), and the A in the fourth position of the tetraloop (A8) were included, while the first A and U in the tetraloop were left unchanged. To vary *K*_fold_, the stem region surrounding the wild-type stem-loop was systematically varied to include 3, 12, or 18 base-pairs; three variants of different GC content (30, 50, or 80%) were designed for each of the longer two stems. Additionally, the directionality of the GC base-pairs that comprise the canonical minimal stem were also systematically flipped to CG to investigate how sensitive MCP binding is to base-pair identity. The entire library was transfected as a plasmid pool into HEK-293T cells either alone or co-transfected with a plasmid coding for MCP. DMS-MaPseq chemical probing was subsequently on each sample and the modified RNAs were sequenced and analyzed using SEISMIC-RNA.^28,29^

All 1,600 library members were recovered after applying stringent quality metrics (**Supplemental Note**) and examined the fraction of sequencing reads that showed DMS-induced mutations at each A or C nucleotide position for each different MS2 sequence. These raw mutation fractions were very low in non-DMS-treated samples, confirming that called mutations are due to DMS modification and not to sequence variability (**Figure S1A**). The mutation fractions between DMS-treated MCP overexpression replicates were highly reproducible (**Figure S1B,** *r* = 0.91). The correlation between the mutation fractions of samples with and without MCP overexpression was significantly lower than between replicates (*r* = 0.74), signifying a difference in measured DMS signal upon MCP overexpression and RNA binding (**Figure S1C**). A normalized difference was computed for each position from the DMS mutation fractions within replicates and with and without MCP overexpression (**Eq. 1**):

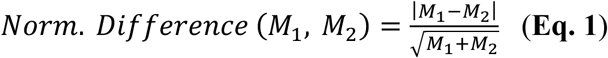

where M1 = mutation fraction of replicate 1 and M2 = mutation fraction of replicate 2.

The distribution of normalized difference values at A and C nucleotides located at positions directly surrounding and including the MS2 stem-loop between MCP overexpression replicates is centered close to 0 with a 95th percentile value of 0.08 (**Figure S1D**). The distribution of normalized difference values at the same positions, but between samples with and without MCP overexpression, is skewed farther to the right with a 95th percentile value of 0.09, indicating that most structure is maintained through MCP overexpression but that MCP binding blocks DMS from reacting with a small number of key bases, increasing the difference between samples (**Figure S1E**).

After confirming that DMS modification was reduced at MCP-contacting positions upon the addition of MCP to the RNA library, the mutational profiles were examined for a library member that we hypothesized should bind MCP strongly in cells. The positional mutation fractions for a wild-type MS2 sequence (**GAGGAUCACCC, long stem, medium GC**) showed the least reactivity (lowest fraction mutated) at bases in the stem and the most reactivity (highest fraction mutated) at positions 5, 7, and 8, or the first, third, and fourth positions of the unpaired tetraloop, which are A, C, and A, respectively, as expected for unpaired positions (**Figure 1D**). This difference in reactivity is consistent with a folded RNA hairpin. The accessibility of the open positions was significantly decreased upon overexpression of MCP (**Figure 1E**). Concordantly, comparing the fraction of reads mutated at each position in the no-MCP condition versus the MCP overexpression condition shows a set of positions that fall below the y=x line (**Figure 1F**), meaning that those positions were more accessible for modification before the addition of protein and are blocked by MCP binding upon its overexpression. The difference in reactivity is consistent with structural data and AlphaFold^40^ predictions (**Figure S2**) where A8 and A2 make hydrogen bonding interactions with Ser47 and Thr45 on the protein on the Watson-Crick-Franklin face, hindering DMS reactivity at these positions. C7 makes π-π stacking interactions with Tyr85 where the Watson-Crick-Franklin face of the base faces the protein interface, blocking it from reacting with DMS. Overall, the differential reactivity at select RNA positions that make direct contact with protein residues indicates that DMS signal can be used to approximate MCP binding.

In contrast, performing the same analyses for a mutant MS2 sequence (**GAG<u>C</u>AU<u>GGG</u>CC**), predicted not to bind MCP, showed a relative lack of disruption to RNA structure at measurable positions (As or Cs), with higher accessibility at the unpaired position 2 (bulge) and 5 (first position in the tetraloop) (**Figure 1G**). However, the mutation fraction of these positions is effectively unchanged upon the addition of MCP (**Figure 1H**), with most positions closely following the y=x line when plotting the no-MCP and MCP overexpression samples against each other (**Figure 1I**). This indicates that, as previously shown, MCP binding is sensitive to MS2 stem-loop mutation and demonstrates that DMS can simultaneously probe RNA folding and RBP binding preferences.

### Stable RNA structures are required for MCP binding

The parameters of the MS2 stem sequence were systematically varied to determine how stem length, GC content, and overall RNA folding contribute to MCP binding of MS2 stem-loops in cells. Changing the stem length (i.e., RNA stability, *K*_fold_) surrounding a wild-type-like MS2 sequence (**GAGGAUUACCC**) confirmed that a three base-pair stem is insufficient for MCP binding, while both 12 and 18 base-pair stems permit hairpin formation and MCP recognition (**Figure 2A-2C**). A long (18 base-pair) stem surrounding the wild-type-like **AUUA** tetraloop produced lower overall mutation fractions for bases in the stem and the first loop position, with increased reactivity at only two positions (A2 and A8), consistent with a folded hairpin structure inaccessible to DMS mutation. The mutation fractions of A2 and A8 both decrease completely to the level of sequencing noise upon MCP addition (**Figure 2A**), consistent with MCP binding to this RNA structure. A medium (12 base-pair) stem shows a similar profile, with several key positions that become protected upon MCP overexpression (**Figure 2B**). In contrast, the shortest stem (three base-pairs) fails to fold sufficiently as indicated by several unpaired bases having a mutation fraction of 0.02 or greater. Consequently, the short stem does not bind MCP, showing a near-perfect correlation between the no-MCP and MCP-overexpression conditions (**Figure 2C**). The GC content of the stem had little effect on MCP binding preferences for the same wild-type stem-loop (**Figure S3A-S3C**), and the stem-length dependencies were consistent across all wild-type or wild-type-like sequences in the library (**Figure S3D**). Mutant MS2 sequences showed smaller changes in response to MCP binding, with largest effects still observed for the medium and long stem lengths (**Figure S3E**).

**Figure 2.**
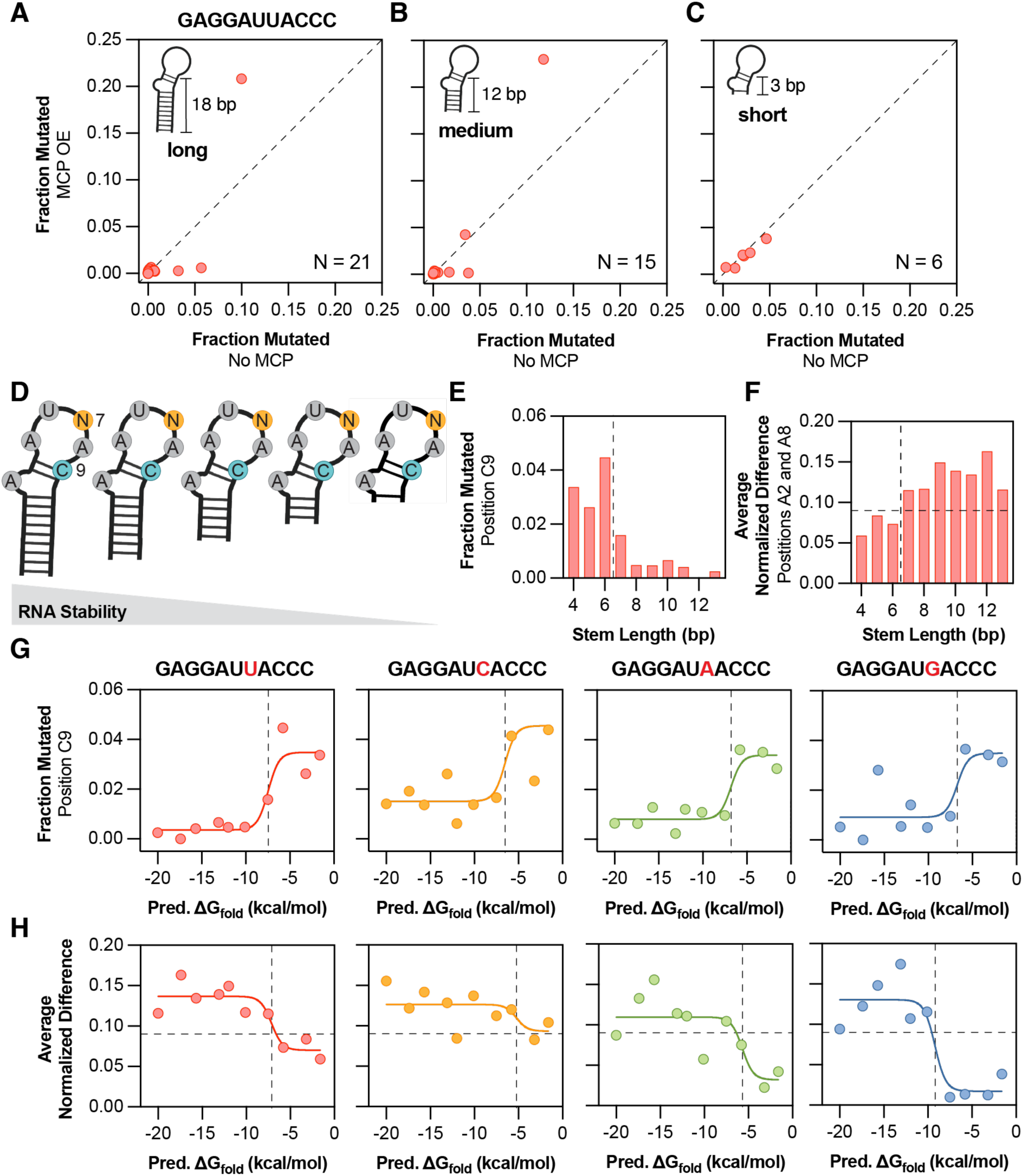
Stable RNA structures are required for MCP binding. (**A-C**) Comparison of DMS-MaPseq reactivity (fraction mutated) at the same positions in the absence (x-axis) and presence (y-axis) of MCP, for the wild-type-like sequence **GAGGAUUACCC** with a long (**A**), medium (**B**), or short (**C**) stem with 80% GC content. Each red circle represents one A or C base (*N*) averaged over two independent replicates. (**D**) A sublibrary of stem-loops with successively increasing stem length, from 4-13 bp, was designed. The position 7 was also varied to each of A, G, C, or U. (**E**) DMS-MaPseq reactivity of a base-paired position, C9 (red bars), for a set of wild-type loop sequences (**GAGGAUUACCC**) with successively longer stems. The vertical dashed lined represented the transition from unfolded hairpins (fraction mutated > 0.02) to folded hairpins (fraction mutated < 0.02). (**F**) MCP binding for the same set of sequences in (**E**), as measured by the average normalized difference to the no-MCP condition at protein-contacting positions 2 and 8 (red bars). The vertical dashed lined represented the transition from unfolded to folded hairpins from (**E**) and the horizontal dashed line represents statistically significant normalized difference (0.09) as established in **Figure S1E**. (**G**) Predicted folding free energies, as calculated by Mfold, versus measured RNA folding by DMS-MaPseq for each of the different tested tetraloop sequences and their varied stem lengths. Left to right, loop sequences with each of U, A, C, and G. Each colored circle represents the fraction mutated of C9 from a sequence of different stem length. The solid line represents the fit to nonlinear regression model relating *P*(unfolded) to Δ*G*_fold_ (see **Methods)**. The dashed vertical line represents the transition from folded to unfolded as calculated by non-linear regression. (**H**) Predicted folding free energy as in (**G**) versus MCP binding as measured by DMS-MaPseq. Left to right, same sequences as in (**G**). Each colored circle represents the average normalized difference of positions A2, A7/C7 (if applicable), and A8. The solid line represents the fit to nonlinear regression model relating *P*(bound) to Δ*G*_fold_ (see **Methods)**. The dashed vertical line represents the transition from bound to unbounded as calculated by non-linear regression and the horizontal dashed line represents statistically significant normalized difference (0.09) as established in **Figure S1E.**

To further probe how MCP binding depends on MS2 stem length, a sublibrary was constructed containing all base substitutions at position 7 within an otherwise wild-type stem-loop (**GAGGAU_ACCC**). For each loop, ten stem mutants were generated by extending the stem one to ten base pairs, for a total of 40 variants (**Figure 2D**). Measuring the DMS mutation fraction in the absence of MCP at the first C past the loop (position C9), which should be base-paired and protected when the RNA folds correctly into a hairpin, revealed that increasing stem length progressively stabilizes RNA folding for a wild-type stem-loop sequence **GAGGAUUACCC** (**Figure 2E**). The longest stems showed the least accessibility, or the most base pairing, with a stem length of around 7 to 8 base pairs required for the stem to fully form (**Figure 2E**). MCP binding was quantified by averaging normalized differences between the +/- MCP conditions at the three loop positions most directly involved in binding (**Eq. 1, Figure S3F**) and defined a binding threshold based on the 95th percentile of these values across the dataset (**Figure S1E**). Both of these metrics suggest that the MCP binding preferences closely follow the stem folding profile, with four additional stem base pairs required for detectable MCP binding to a wild-type stem loop (**Figure 2F**).

These analyses were expanded by using the mFold RNA prediction software^41^ to calculate the predicted folding free energy (Δ*G*_fold_) for each of the 4 loop sequences across their 10 stem lengths. DMS mutation fractions in the absence of MCP at position C9 scaled logarithmically with predicted Δ*G*_fold_ (**Figure 2G**): below approximately -5 kcal/mol, DMS reactivity at position C9 was nearly zero, indicating complete stem pairing. Above this value, DMS accessibility sharply increased before reaching a plateau, consistent with unfolded or unstable RNA stems (**Figure 2G**). It is likely that the RNA hairpin structures are largely destabilized in cells relative to *in vitro* computational predictions,^22,42,43^ which contributes to the hypersensitive relationship between predicted structure and measured DMS reactivity. Comparing predicted Δ*G*_fold_ to average normalized differences (MCP binding) across the four loop families revealed the same trend, with MCP binding to each loop occurring only when Δ*G*_fold_ for the sequence is predicted to be below -5 kcal/mol (**Figure 2H**). For the wild-type sequence **GAGGAUUACC**, MCP binding decreased sharply at the same stem length where folding was lost (**Figure 2G, 2H; red**), indicating a direct link between RNA structure and protein recognition. The other wild-type-like sequence **GAGGAUCACCC** showed similar RNA folding behavior, maintained binding across all stem lengths (**Figure 2G, 2H; orange**), suggesting that the AUCA loop sequence has higher intrinsic affinity for MCP and requires less structural stabilization for recognition. The other two loop sequences (**GAGGAU<u>A</u>ACCC** and **GAGGAU<u>G</u>ACCC**) required significantly more stable RNA structures, with Δ*G*_fold_ less than -10 kcal/mol, before MCP binding was detected, although both sequences folded at the same -5 kcal/mol as the other two variants (**Figure 2G, 2H; green and blue**). This indicates a slight disfavoring of MCP recognition of the non-wild-type AUAA and AUGA loop sequences such that a more stable RNA structure is required for MCP to bind.

### Loop sequence is the second-biggest determinant of MCP binding preferences

After determining the importance of stable RNA structures for strong MCP-MS2 recognition, we examined the contribution of individual nucleotides to binding specificity. A wild-type-like MS2 sequence (**GAGGAUUACCC, long stem, medium GC**) was compared to all possible single-nucleotide substitutions at positions 2, 5, 7, and 8 (**Figure 3A**). This systematic set of variants allows us to link single-base changes to their effects on MCP binding; for example, how mutating position 2 from A to U, C, or G alters binding.

**Figure 3.**
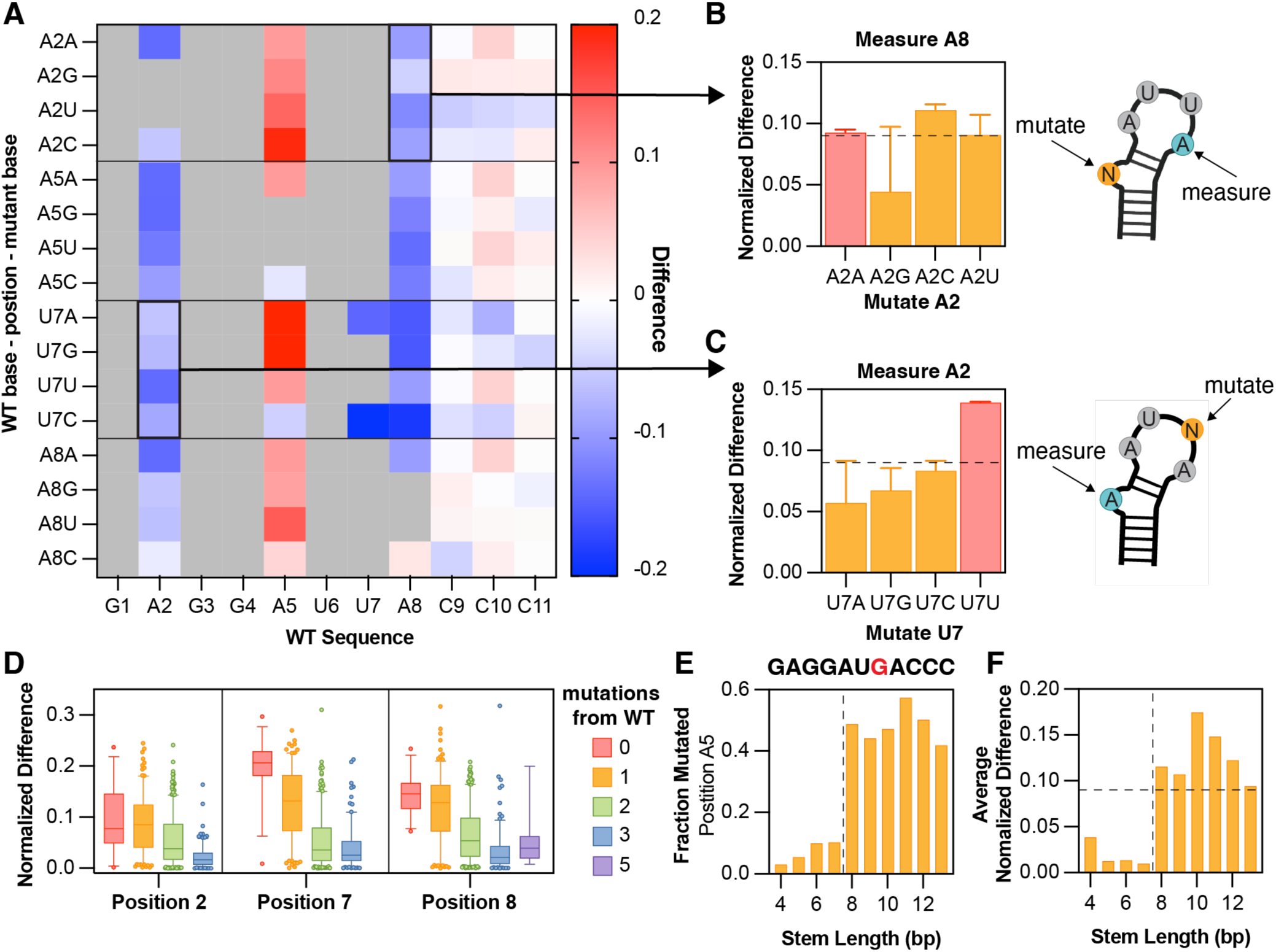
Loop sequence is the second-biggest determinant of MCP binding preferences. (**A**) Heatmap of the normalized difference in DMS-MaPseq signal from MCP overexpression to MCP absence at each stem-loop position (x-axis) in response to each mutation elsewhere in the sequence (y-axis). Grey boxes, non-measured positions (Gs or Us) or mutations that were not tested. (**B**) Quantification of the normalized difference in DMS-MaPseq signal upon MCP overexpression at position A8 in response to different base mutations made at the position 2. Each colored bar was averaged over two independent replicates. Error bars indicate the range. (**C**) Quantification of the normalized difference in DMS-MaPseq signal upon MCP overexpression at position A2 in response to different base mutations made at position 7. Each colored bar was averaged over two independent replicates. Error bars indicate the range. (**D**) Normalized difference in DMS-MapSeq signal at positions 2, 7, and 8. Boxes show first to third quartiles, whiskers show down to the 5^th^ and up to the 95^th^ percentile, and outliers are shown as circles. Colors indicate the number of mutations from the wild-type sequence (zero = red; one = orange; two = green; three = blue; and five = purple). (**E**) DMS-MaPseq reactivity (orange bars) of position A5 for a set of loop sequences (**GAGGAU<u>G</u>ACCC**) with successively longer stems. The vertical dashed lined represented the transition from unbound hairpins to bound hairpins as calculated in Figure 2G. (**F**) Average normalized difference to no MCP for positions A2 and A8 for the same set of sequences in (**E**). The vertical dashed lined represented the transition from unbound to bound hairpins from (**E**) and the horizontal dashed line represents statistically significant normalized difference (0.09) as established in **Figure S1E**.

Because MS2 sequences mutated at the bulge nucleotide (position 2) are known to be permissive to MCP binding, variation at this position provides a useful test case for our approach. Although DMS reactivity cannot be measured directly for G or U at this position (since they lack DMS-reactive sites on the Watson-Crick-Franklin face), other protein-contacting bases in the sequence can serve as proxies for how binding specificity is altered upon mutation. Position A8, which directly contacts the MCP dimer, was used as a constant “watch” position. Normalized differences at A8 showed that all bases except G are tolerated at position 2 (**Figure 3B**), with G being heavily disfavored for binding. This is likely because a G bulge can form unintended base pairs with opposing stem Cs, disrupting proper MS2 folding and occluding MCP access to the exposed bulged nucleotide.

A similar strategy was applied to position 7, using A2 as the “watch” position. Here, the wild-type-like C is strongly preferred at position 7 (**Figure 3C**), with all other bases being slightly disfavored. In contrast, using the A5 as the “watch” position shows the opposite trend, with large increases in accessibility upon MCP binding reflected by positive difference values in response to mutations at either positions 2 or 7 (**Figure 3A**). This suggests a conformational change in the MS2 loop upon MCP binding, potentially involving increased accessibility of the Watson-Crick-Franklin face or increased nucleophilicity of the N1 position of adenosine in the bound state (**Figure S4**), which can serve as an additional proxy for MCP binding. Notably, the largest increases in reactivity at A5 occur in response to mutations that substitute purines for pyrimidines or vice versa (**Figure 3A**), suggesting that base identity influences both the conformation of the tetraloop in the bound state and its reactivity to DMS.

Together, these measurements demonstrate that comparing position-specific differences in DMS accessibility with and without MCP enables quantitative measure of RNA-protein interaction specificity. Among all variables tested, sequence similarity to the wild-type MS2 loop is the strongest determinant of MCP binding at the three positions that directly contact the protein: even a single mutation markedly reduces the normalized difference between the +/- MCP conditions (**Figure 3D, Figure S4A**). Sequences with two or more mutations away from wild type show little detectable binding, underscoring the high specificity of both the MS2 hairpin structure and its interaction with MCP (**Figure 2H**).

By contrast, stem length and stem GC content have much smaller effects on binding. Increasing stem length beyond 12 base pairs does not significantly alter binding at any of the three positions (**Figure S4B–D, pink and orange boxes**), although constructs with very short stems (3 bp; **Figure S4B–D, green boxes**) fail to bind MCP efficiently, consistent with the results in **Figure 2**. Similarly, variation in stem GC content does not further affect binding beyond the effects caused by altering the loop sequence (**Figure S4E–G**). Finally, normalized difference values at other stem-loop positions reported not to directly interact with MCP remain largely unchanged across increasing mutational distances from wild type (**Figure S4H**), confirming that positions 2, 7, and 8 are the primary reporters of MCP binding.

Lastly, the increased accessibility at the first position of the tetraloop (A5) observed for pyrimidine to purine substitutions in **Figure 3A** was examined in more detail. Using the stem-loop sequence **GAGGAU<u>G</u>ACCC**, the DMS mutation fraction was measured directly in the presence of MCP, rather than the normalized difference to no MCP, and found a pronounced increase in A5 reactivity upon the addition of MCP (**Figure 3E**) that coincides with the onset of MCP binding to an eight base-pair stem (**Figure 3F**). This increase is not observed for the wild-type **GAGGAUUACCC** sequence, despite measurable binding at a similar stem length (**Figure 2F, Figure S6C**). Although the precise mechanism underlying the increased A5 reactivity remains unclear, these results highlight the power of high-dimensional DMS datasets to reveal RNA-protein interaction dynamics across every nucleotide position, not only at those directly contacting the protein.

### DMS-MaPseq can quantify RBP binding across a wide range of concentrations

A system in which protein levels could be controlled over a wide range was next examined to evaluate the sensitivity of DMS-MaPseq for measuring binding specificity. Measuring binding at lower protein concentrations is an important test of this approach because most endogenous RBPs exist at subsaturating concentrations with respect to their RNA targets.^42,44,45^ The same MCP dimer construct was fused to an inducible DHFR degron, a constitutive protein degradation tag that can be stabilized by increasing concentrations of trimethoprim (TMP) (**Figure 4A**).^46,47^ Increasing TMP from 0 to 10 µM monotonically raised MCP levels (**Figure S5A**), although these levels remained well below the degronless MCP. While some residual MCP may persist at 0 TMP due to DHFR leakiness, the large differences between TMP-controlled and overexpressed concentrations allow measurements over a wide range of protein concentrations, including in a significantly subsaturating regime.

**Figure 4.**
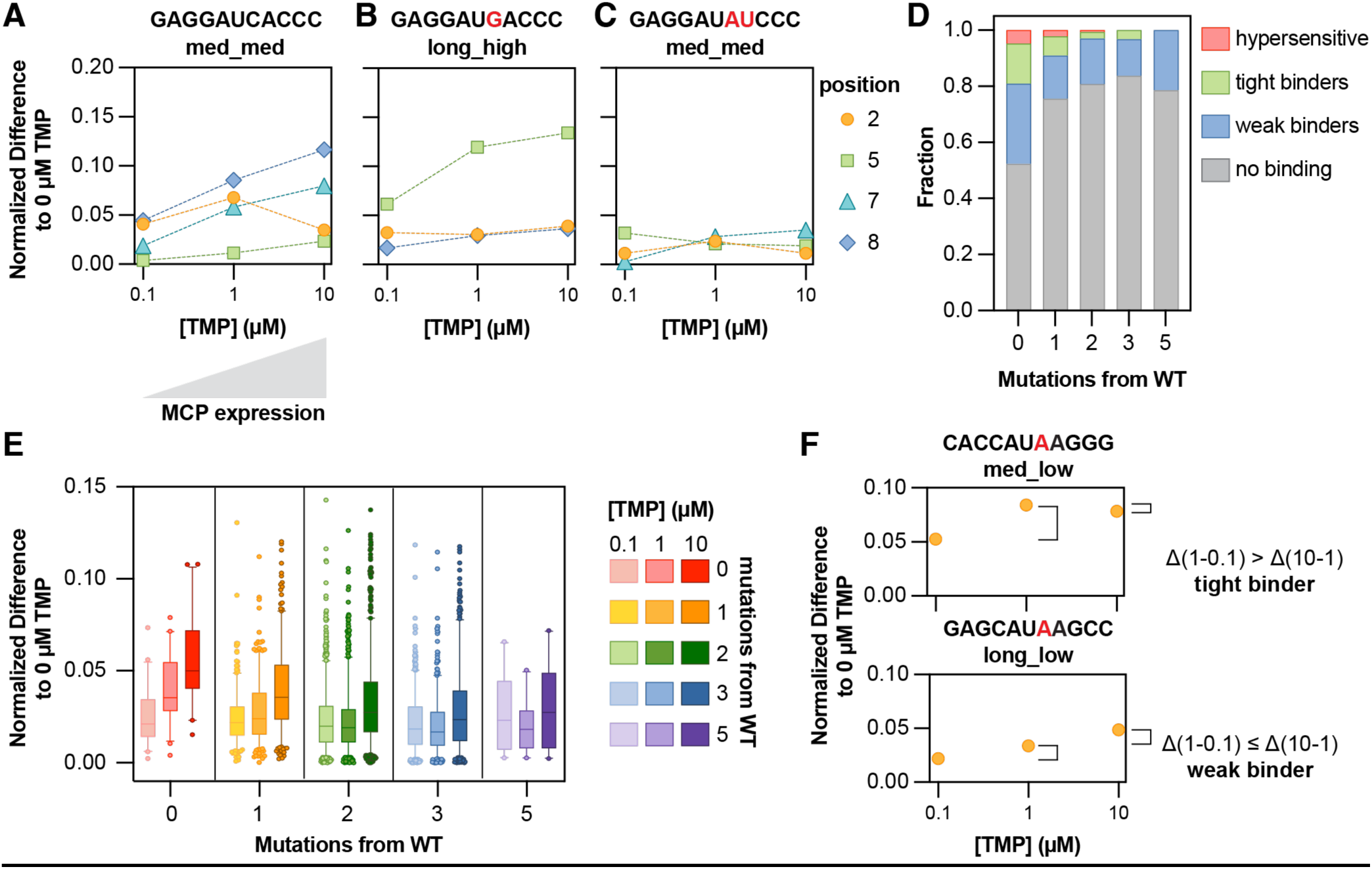
DMS-MaPseq can quantify RBP binding across a wide range of concentrations. (**A-C**) TMP dose-response binding curves for (**A**) A wild-type MS2 sequence, (**B**) A sequence with one C to G mutation at position 7, and (**C**) a two-mutation, non-binding sequence (right). Each dashed line represents a different measured RNA nucleotide position that makes contact with MCP: position 2 (orange), position 5 (green), position 7 (turquoise), and position 8 (blue). X-axis, TMP dose; y-axis, normalized difference at the given position and TMP dose to the 0 TMP condition. (**D**) The fraction of sequences with increasing numbers of mutations away from wild-type MS2 classified as non-binding, weak, tight, or ‘hypersensitive’ (large binding differences from 0.1 to 1 and from 1 to 10 μM TMP). The threshold for any binding activity between two given concentrations is set using the 95th percentile of the distribution of all 0.1 to 1 μM differences. (**E**) Average responsiveness to TMP for sequences with increasing number of mutations away from wild-type MS2, as measured by the average normalized difference to 0 TMP at the three positions (2, 7, 8). Boxes show first to third quartiles, whiskers show down to the 5^th^ and up to the 95^th^ percentile, and outliers are shown as circles. Only measurements with normalized difference less than 0.15 are shown. Colors indicate the number of mutations from the wild-type sequence (zero = red; one = orange; two = green; three = blue; and five = purple). Shading indicates the TMP dose (light = 0.1 μM; medium = 1 μM; and dark = 10 μM). (**F**) Dose responsiveness to TMP as measured by averaged normalized difference to 0 TMP for two example sequences. Top, classified as a ‘tight binder’, where the difference between its average binding at 1 μM TMP and 0.1 μM TMP is bigger than the difference between 1 and 10 μM TMP (saturating at lower TMP dose). Bottom, classified as a ‘weak binder’, where the binding difference from 1 to 10 μM TMP is bigger than 0 to 0.1 TMP (sub-saturating even at maximum tested TMP dose).

We calculated the normalized difference between the DMS mutation fractions at each position between each TMP dose and the 0 TMP condition. For a wild-type sequence **GAGGAUCACCC,** normalized differences increased monotonically at positions 7 and 8 with increasing TMP, reflecting non-saturating binding as MCP concentration increased (**Figure 4B**). In contrast, after mutation at those positions (**GAGGAU<u>AU</u>CCC**), very little response to TMP was measured, indicating an overall non-binding sequence (**Figure 4C**). When only position 7 was mutated to G (**GAGGAU<u>G</u>ACCC**), there was much less response to TMP increase except for at position 5 (**Figure 4D**), consistent with the purine-driven accessibility increase seen in **Figure 3F**. At subsaturating MCP concentrations, MCP appears sufficient to induce a concentration-dependent conformational change at position A5 of the tetraloop but does not make measurable contacts with the A2 or A8. This might suggest that the bound conformation is different from that of the wild-type or wild-type-like MS2 sequences.

Overall, the average normalized difference for each sequence compared to the 0 TMP condition generally increased with TMP dose. Wild-type sequences and those with a single loop mutation showed the strongest response and highest overall binding (**Figure 4E, Figure S5B**). Sequences with three or five mutations exhibited minimal binding but still showed modest TMP responsiveness, indicating that varying MCP concentration can reveal sub-threshold interactions that might otherwise be overlooked in bulk overexpression experiments.

To further quantify this, the responsiveness of each sequence was calculated by comparing average binding between TMP doses of 0.1 and 1 µM and between 1 and 10 µM, defining a threshold from the 95th percentile of the 0.1–1 µM differences across the dataset (**Figure S5C**). Sequences above the threshold at 0.1-1 µM but below it at 1-10 µM were classified as sensitive, or ‘tight’ binders, saturating at low TMP and thus exhibiting a lower effective cellular *K*_d_ (**Figure 4F, top**). Sequences below the threshold at low TMP but above it at high TMP were classified as less sensitive, or ‘weak’ binders (**Figure 4F, bottom**). Notably, both sequences shown in **Figure 4F** were well above the binding threshold when incubated with degronless MCP (binding metrics of 0.2, top, and 0.16 bottom; threshold 0.09 (**Figure 2**)). However, modulation of MCP concentration effectively revealed variabilities in their affinities in cells and demonstrated the need for a sensitive measurement technique to identify sub-saturating binding dynamics.

The entire library was classified into four binding groups: zero binding, weak binding, tight binding, and hypersensitive (above the responsiveness threshold in both concentrations), and saw that only sequences with fewer than 2 mutations from the wild-type sequence were ever classified as tight binders or hypersensitive (**Figure 4G**). As mutational distance increased, the fraction of weak binders decreased while non-binders increased. These classifications agreed with data using the degronless MCP, with tight and hypersensitive sequences showing significantly higher normalized binding in the highly-expressed degronless MCP dataset (**Figure S5D**). Together, these results show that tuning RBP abundance reveals affinity-dependent binding and conformational responses that are obscured under saturating conditions, underscoring the power of DMS-MaPseq to capture quantitative binding dynamics in cells.

### Quantitative framework for estimating fraction bound from MCP overexpression

Finally, we established a quantitative framework that converts the DMS-MaPseq signal for each RNA into an estimate of the fraction of RNAs bound by MCP inside cells. At each RNA position *j*, the measured mutation fraction reflects the sum of the probabilities (*P*) of unfolded, folded, and bound RNA states and the DMS reaction rate of that state (*k*_DMS_, **Eq. 2, Methods**). The observed mutation fraction in each experiment for a given position is thus:

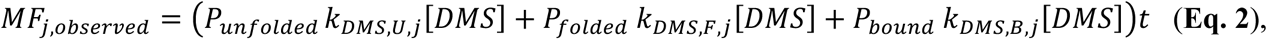

where k*_DMS, U/F/B, j_* is the rate of modification (per unit of DMS concentration, denoted by [DMS]) at position j when the RNA is in the unfolded, folded and unbound, or bound states respectively.

Using the stem length and folding analyses in **Figure 2**, we have determined that every library member with a medium or long stem was determined to be folded in cells, evidenced by a lack of DMS reactivity at the paired stem bases. By restricting our analyses to only those library members, we know that the fraction of unfolded RNAs is negligible (*P_unfolded_* ≈ 0). Additionally, only the RNA positions that directly contact MCP to approximate binding were used for this analysis, for which the rate of DMS reactivity (*k*_DMS,B_) is essentially zero when bound to MCP. Under these constraints, the observed mutation fraction is proportional to the fraction of molecules that are folded but unbound at that position.

Thus, for each position *j*:

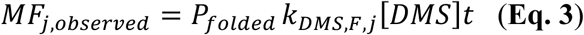

When the sequence is completely bound by MCP, *P_bound_* = 1 and *P_folded_* = 0; inversely, *P_folded_* = 1 indicates a complete lack of binding at that position.

For each mutant sequence, a relative *P_folded_* was computed by comparing its mutation fraction to the wild-type at the same base. Importantly, this analysis was restricted to only common bases shared between the reference wild-type and each mutant sequence to ensure a constant rate of DMS modification, which is variable per base. All measured positions in the absence of MCP can be represented by *P_folded_* = 1. This leads to the following equation for the differential probability of being folded (but not bound) between a mutant RNA compared to a WT sequence, based on the observed mutation fractions of each with and without MCP (see **Methods** for full derivation):

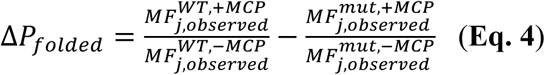

for each of the shared bases between the WT reference and each library member. The average was taken for each of those positions, which leaves us with a simplified *ΔP_folded_*, or fraction folded, for each sequence. Since these RNAs were assumed to be entirely folded, the fraction bound is simply:

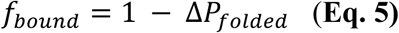

For this dataset, the wild-type reference sequence **GAGGAUCACCC, long stem, medium GC**, shown in **Figure 1**, was assumed to be completely bound: the DMS mutation fractions at positions 2, 7, and 8 decrease approximately to 0 upon the addition of MCP (**Figure 1D-F**), enabling calculation of an absolute fraction bound. When applying this framework to wild-type and mutant RNA sequences for which the wild-type RNA is not fully bound, *f_bound_* should instead be reported as *ΔP_bound_,* representing occupancy relative to the wild-type sequence.

This analysis provides a direct, quantitative measurement of the fraction of each RNA species that is bound in living cells. As expected, the fraction bound is very high (approaching 1) for each wild-type-like sequence in the library and decreases as the number of mutations away from the wild-type sequence increases (**Figure 5A**). Additionally, the fraction bound is well-correlated with the qualitative normalized difference used in earlier analyses to rank binding strengths of different RNA sequences (**Figure S6A**). Collapsing measurements across RNAs with the same stem-loop but different stem lengths or GC content reveals a clear mutational walk, as progressively diverging from the wild-type loop reduces the fraction bound in a near-monotonic fashion (**Figure 5B**). These data allow an approximate percentage of how much each base contributes to overall binding. For example, a single mutation in the third tetraloop position from C to A (**GAGGAU<u>A</u>ACCC**) decreases the fraction bound only by ∼5-10% (**Figure 5B, third bar from left**), but a single mutation in the fourth tetraloop position from A to C (**GAGGAUC<u>C</u>CCC**) decreases binding by ∼20% (**Figure 5B, fifth bar from left**).

**Figure 5.**
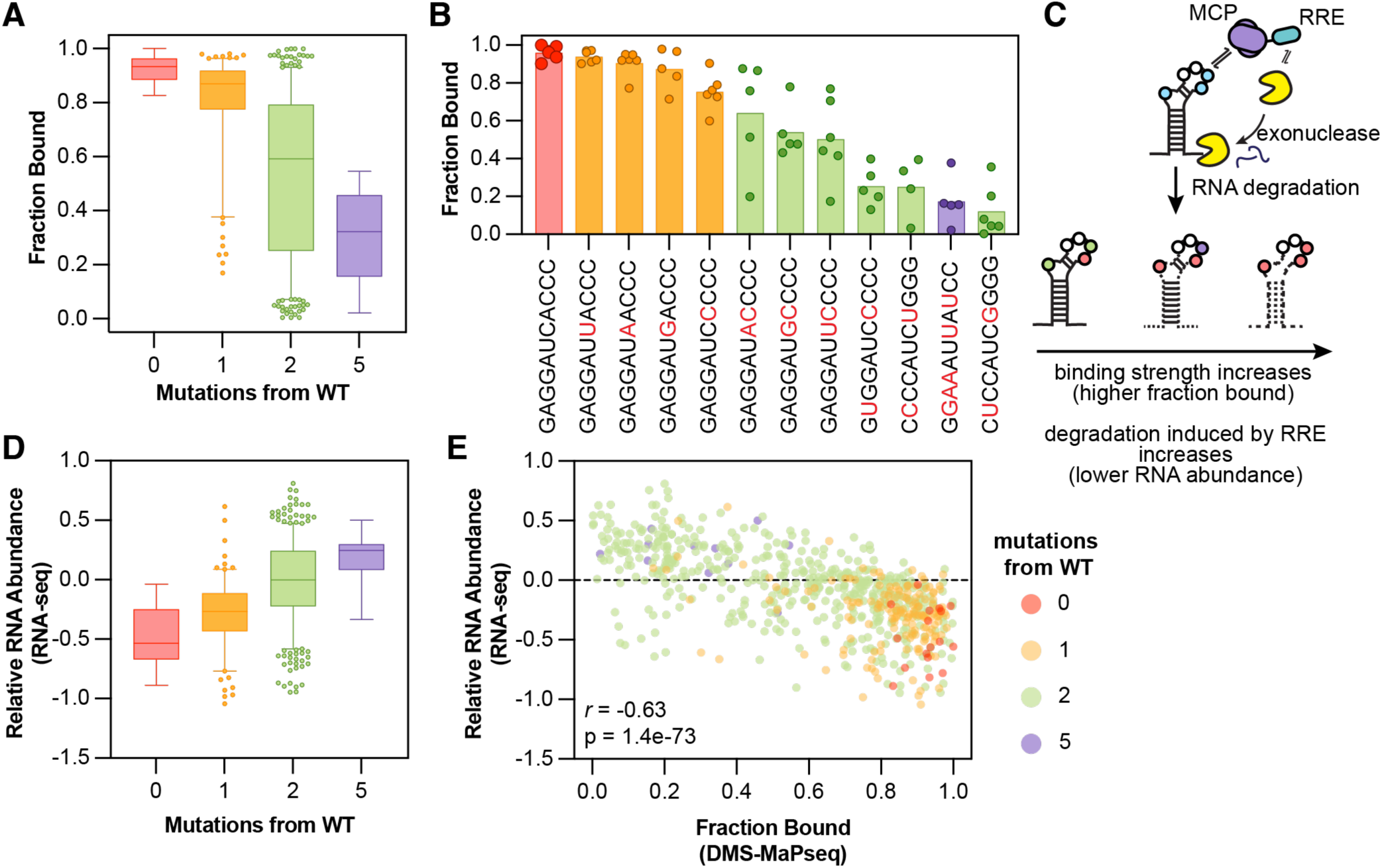
DMS-MaPseq predicts RBP occupancy and downstream gene regulation in living cells. (**A**) Measurements of the fraction of RNA molecules bound by MCP for library members with increasing numbers of mutations away from the MS2 consensus. Boxes show first to third quartiles, whiskers show down to the 5^th^ and up to the 95^th^ percentile, and outliers are shown as circles. Colors indicate the number of mutations from the wild-type sequence (zero = red; one = orange; two = green; and five = purple). Only measurements with fractions bound less than 1 are shown. (**B**) Measurements of fraction bound for example loop sequences. Circles show measurements for each loop with different stem lengths (medium, long) or GC contents (low, medium, high); bars show the average of those measurements. Bar colors show the number of mutations in each loop as compared to wild-type as in (**A**). (**C**) Schematic of RNA degradation experiment. An RNA-downregulatory effector domain (RRE) from the protein NANOS1 (amino acids 21-100) known to induce degradation of bound transcripts is fused to MCP, which is overexpressed in the presence of the same MS2 stem-loop library. As MCP occupancy increases, as measured by DMS-MaPseq, RNA abundance of a given library member should decrease as degradation is more effective. (**D**) Relative abundances of MS2 stem-loop library members, measured as a log_2_FoldChange between overexpression of the MCP-RRE construct compared to overexpression of RRE alone. Sequences are separated by increasing numbers of mutations away from the MS2 consensus as in (**A**). (**E**) Comparison between the predicted fraction bound of each stem-loop library member from DMS-MaPseq measurements and its measured relative abundance upon MCP-RRE overexpression. Colors represent mutational distance from the wild-type MS2, n = 667 sequences. Pearson r = -0.63; p-value = 1.4 *10^-73^.

### Functional Consequences of MCP Specificity

To test whether the quantitative measurements of MCP-RNA binding could predict functional consequences inside cells, we fused MCP to a potent RNA-regulatory effector domain (RRE) from the protein NANOS1 known to trigger degradation of its bound RNA targets (**Figure 5C**).^46^ The MCP-RRE fusion was overexpressed together with the same stem-loop library and quantified the steady-state relative abundance of each RNA variant by RNA-seq after 48 hours compared to an MCP-alone control (**Figure S6B**). As expected, wild-type-like stem-loops, which were determined to be almost fully bound, showed the strongest depletion. RNA abundance increased progressively as the stem–loop accumulated more mutations (**Figure 5D**), demonstrating that stem-loop sequence is a strong determinant of RNA fate.

The inferred fraction bound from DMS-MaPseq measurements directly and quantitatively predicted the degree of RNA depletion. Across all library members, an anticorrelation was observed between fraction bound and RNA abundance (*r* = –0.63; **Figure 5E**). Thus, the binding measurements in **Figure 5A** and **5B** are not merely descriptive: they predict RNA lifetimes in cells, linking the physical occupancy of an RBP to the magnitude of its regulatory output.

## CONCLUSIONS

The development of this quantitative DMS-MaPseq framework provides an experimental platform for dissecting RNA-protein interactions at single-nucleotide resolution directly within living cells. By integrating RNA-centric metrics into RBP binding analyses, this system enables systematic probing of the functional consequences of cell state changes and mutational effects that alter both RNA structure and protein affinity in native contexts. This framework also expands the possibilities for creating tunable tools or therapeutics whose RNA-binding properties can be predictably modulated by specific structural modifications or sequence changes.

Using the well-studied MCP-MS2 system as a test case, a library of systematically mutated MS2 stem-loops was generated to map how individual nucleotide identities and RNA structures contribute to MCP binding specificity. These data show that single-nucleotide changes within the MS2 loop can substantially alter MCP binding, and that stem stability strongly influences the ability of the loop to adapt the correct conformation for recognition by MCP. Importantly, these data imply that stable RNA structures must form in cells before RBP binding, indicating that sequence recognition alone is often insufficient for specific recognition. This may explain why RBP motifs are often highly degenerate^48,49^ and why binding patterns are difficult to predict without transcriptome-wide pulldown data: sequence prediction models do not capture the substantial contribution of RNA structure to binding affinities and probabilities^50^.

The TMP-inducible MCP system demonstrates that DMS-MaPseq is sensitive for binding detection across a wide range of varying protein concentrations, including low levels. This opens opportunities to investigate the interplay between protein concentration, binding affinity, and cellular dynamics. Shorter DMS treatments, or time-course experiments, could reveal transient or weaker interactions that might be missed under overexpression conditions, offering a method to probe the full spectrum of RNA-protein affinities. Additionally, while this study focuses on a single RBP and its cognate stem-loop, this sensitivity indicates that the method is readily extendable to endogenous RNA-protein interactions and more complex networks. Applying the same DMS-MaPseq workflow *in vitro* and comparing it to in-cell measurements could benchmark predictions of RNA structure and RBP binding, account for cellular context effects, and refine models of RBP specificity. This is particularly valuable for understanding interactions in crowded, structured, or highly dynamic RNA environments where *in vitro* affinities may not fully recapitulate behavior in cells^51^. DMS-MaPseq could also be combined with complementary approaches to measure RBP binding, such as UV crosslinking to provide orthogonal confirmation of binding events. While DMS reports on structural accessibility and indirect occupancy, UV crosslinking captures direct contact between RNA and protein^52^, allowing for comparisons of transient and stable interactions. Integrating these methods could yield a more complete map of RBP-RNA interactions.

The quantitative framework produced here allows us to link the fraction of RNA bound by MCP directly to functional outcomes in cells. Using the MCP-RRE fusion experiments, hairpins with higher fractional occupancy experienced stronger RNA depletion, while those with reduced binding remained more abundant. This establishes that DMS-MaPseq-derived occupancy measurements are not merely descriptive, but predictive of regulatory output, allowing for a predictive functional model of RNA sequence and structural changes in cells. In principle, this approach could be extended to other RBPs or RNA effectors to predict functional outcomes such as splicing efficiency, translation, or decay, based solely on measured occupancy. This approach also provides a new strategy for engineering tunable RNA tools: MS2 hairpins can now be designed to achieve graded MCP affinities by varying stem length, GC content, or specific loop nucleotides. Such synthetic RNA hairpins could then be used to fine-tune regulatory circuits, generate dose-dependent RNA-protein interactions, or develop RNA-based therapeutics whose function depends on predictable binding dynamics.

## SUPPORTING INFORMATION

Supporting information: Figures S1-S6, containing figures of internal experimental controls, AlphaFold structural depictions, and additional analyses to support claims made in main text. Methods section, containing all experimental details.

## ACKNOWLEDGEMENTS

We thank Maya Schauber for experimental assistance, Casper L’Esperance-Kerckhoff for help generating supplemental datasets, and Yaara Finkel and all members of the Hagler lab for helpful conversations. This work was supported by NIH-NCI F30CA287739-01 (A.R.T.), a Stanford Bio-X Bowes Fellowship (A.R.T.), a Stanford Sarafan Chem-H Chemistry-Biology Interface Fellowship (A.R.T.), NIH-NIGMS MIRA R35GM12894701 (L.B.), Howard Hughes Medical Institute Hanna H. Gray Fellowship (L.D.H), and CPRIT RR240063 (L.D.H.)

## AUTHOR CONTRIBUTIONS

A.R.T. and L.D.H. conceptualized and designed the study. A.R.T. performed all DMS structural profiling experiments with assistance from T.H.N. A.R.T. performed and analyzed RNA-seq experiments. A.R.T. and L.D.H. analyzed and visualized the data with assistance from M.M. J.D.T.C. performed AlphaFold structural predictions and molecular dynamics simulations. A.R.T. and L.D.H. wrote the fraction occupancy calculations. A.R.T. and L.D.H. wrote the manuscript with input from L.B. and contributions for all authors. L.D.H. supervised the study.

---

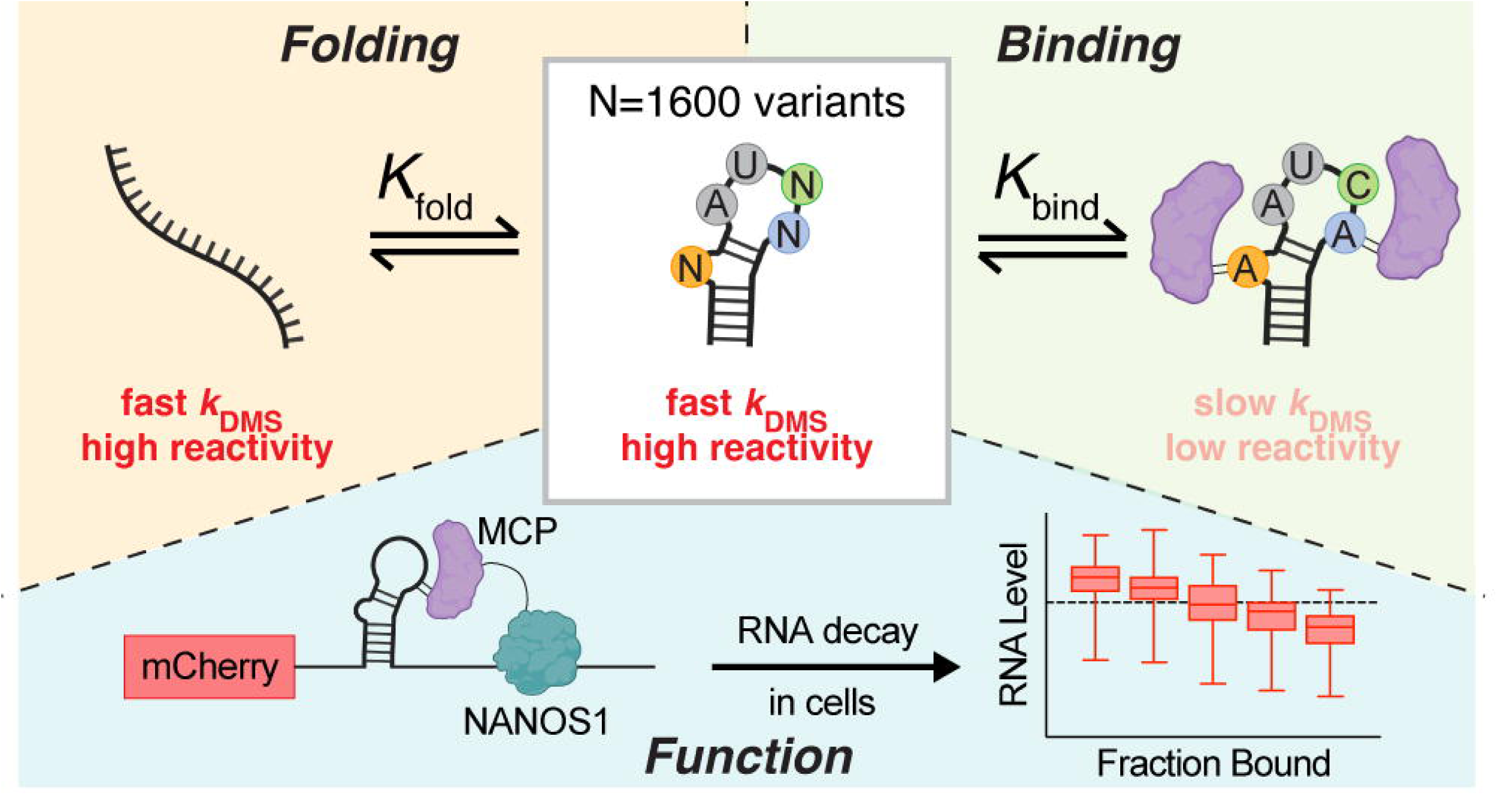

