## Supplementary Information for "Structural probing of RNA hairpins quantifies protein occupancy on RNA and links it to function in human cells"

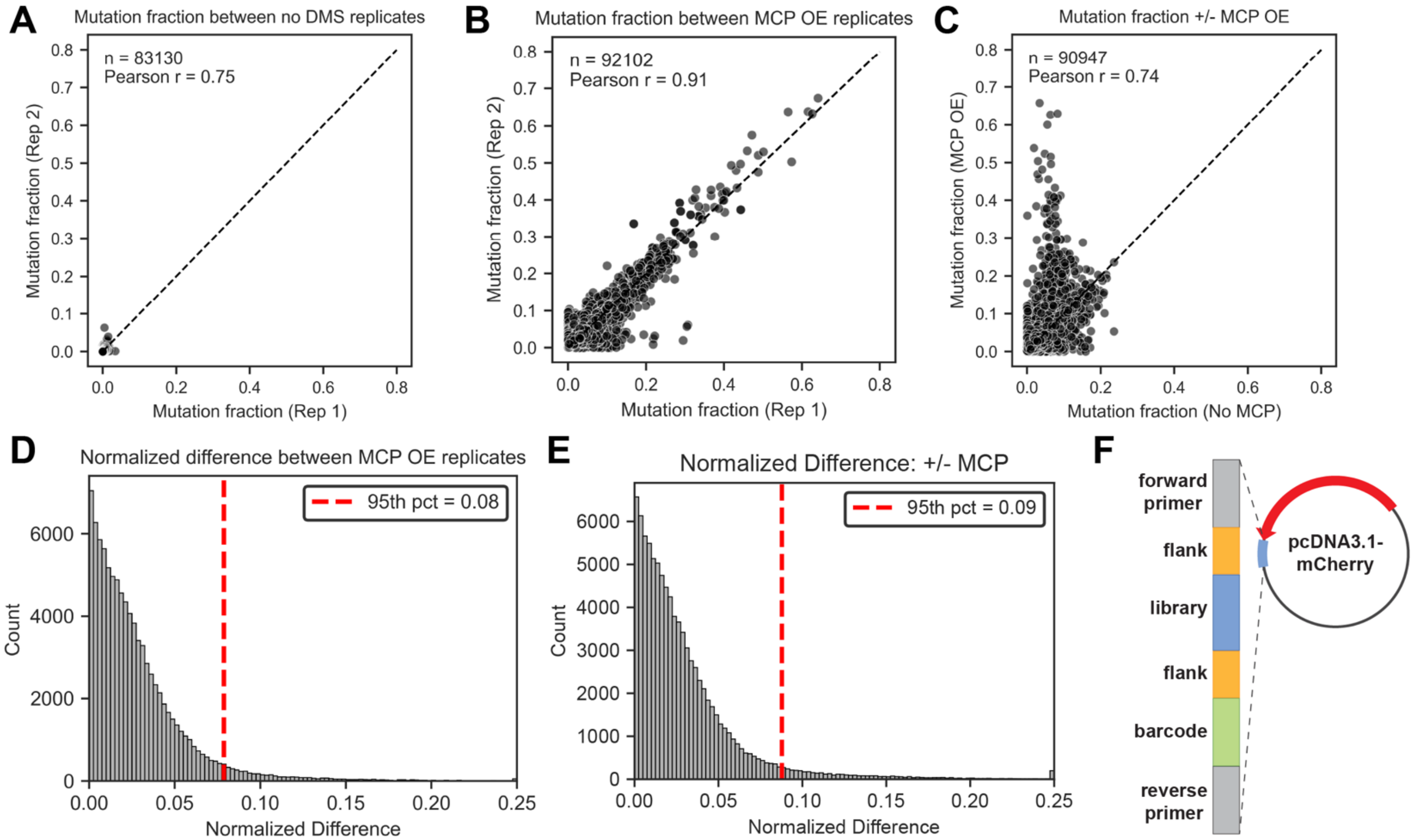


**Figure S1. DMS-MaPseq replicates and metrics, related to Figure 1.** (A) Mutation fractions as measured by DMS-MaPseq for all A and C bases in the MS2 library from samples transfected into HEK293T cells and not treated with DMS. X-axis, replicate 1; y-axis, replicate 2; N = 83,130 positions measured. (B) Mutation fractions as measured by DMS-MaPseq for all A and C bases in the MS2 library from samples transfected into HEK293T cells with overexpression of MCP, treated with DMS. X-axis, replicate 1; y-axis, replicate 2; N = 92,102 positions measured. (C) Comparison of mutation fractions, averaged over replicates, for A and C bases in the MS2 library from samples transfected into HEK293T cells with (y-axis) and without (x-axis) MCP overexpression, treated with DMS. N = 90,947 unique positions measured. (D) Distribution of normalized difference values between two replicate samples of A and C bases in the MS2 library transfected into HEK293T cells with overexpression of MCP, calculated using Eq. 1. X-axis, normalized difference value; vertical line, 95th percentile value and significance cutoff. (E) Distribution of normalized difference values between samples of A and C bases in the MS2 library transfected into HEK293T cells with and without overexpression of MCP, calculated using Eq. 1. X-axis, normalized difference value; vertical line, 95th percentile value and significance cutoff. (F) Library design for the MS2 hairpin library.

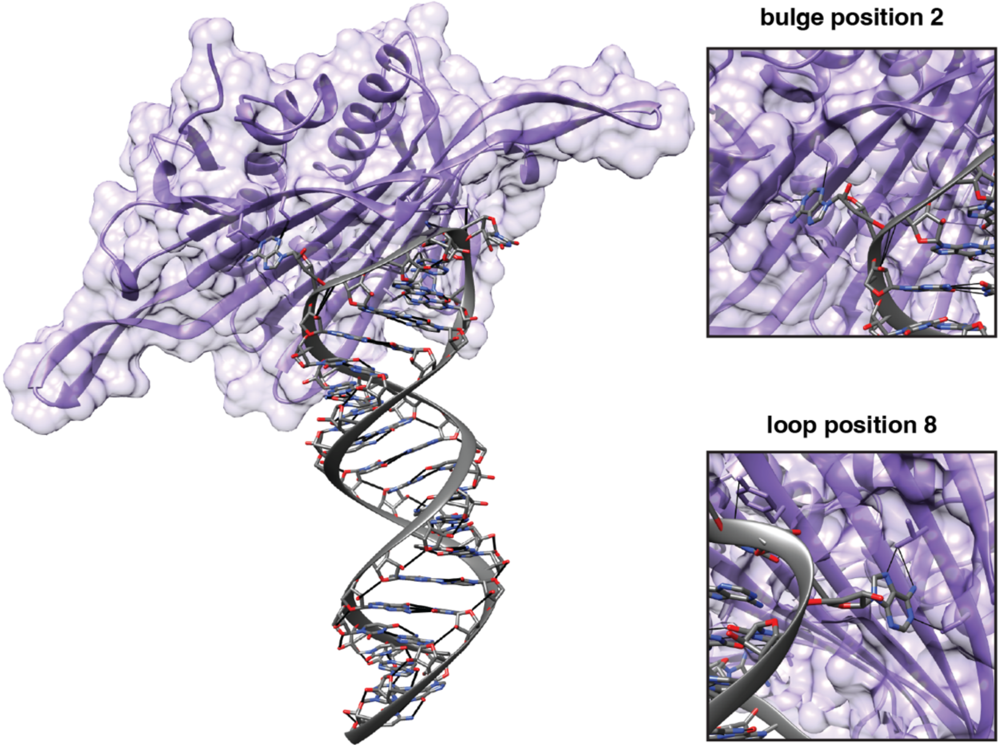


**Figure S2. AlphaFold Prediction of RNA-protein complex**. The structure of GAGGAUCACCC_med_med was predicted with two copies of MCP (Uniprot: P03612) using the AlphaFold server.[1] The resulting complex was visualized and analyzed in Chimera.[2] (left) The protein, MCP, is shown in purple. The RNA bases are shown in grey. The black lines indicated predicted hydrogen bonds based on the angle and distance of the hydrogen bond donor and acceptor. (top right) Hydrogen bonding from threonine 45 to the N1 position of A2 and from serine 47 to the N3 position of A2. (bottom right) Hydrogen bonding from threonine 45 to the N6 and N7 positions of A8 and from serine 47 to the N1 position of A8.

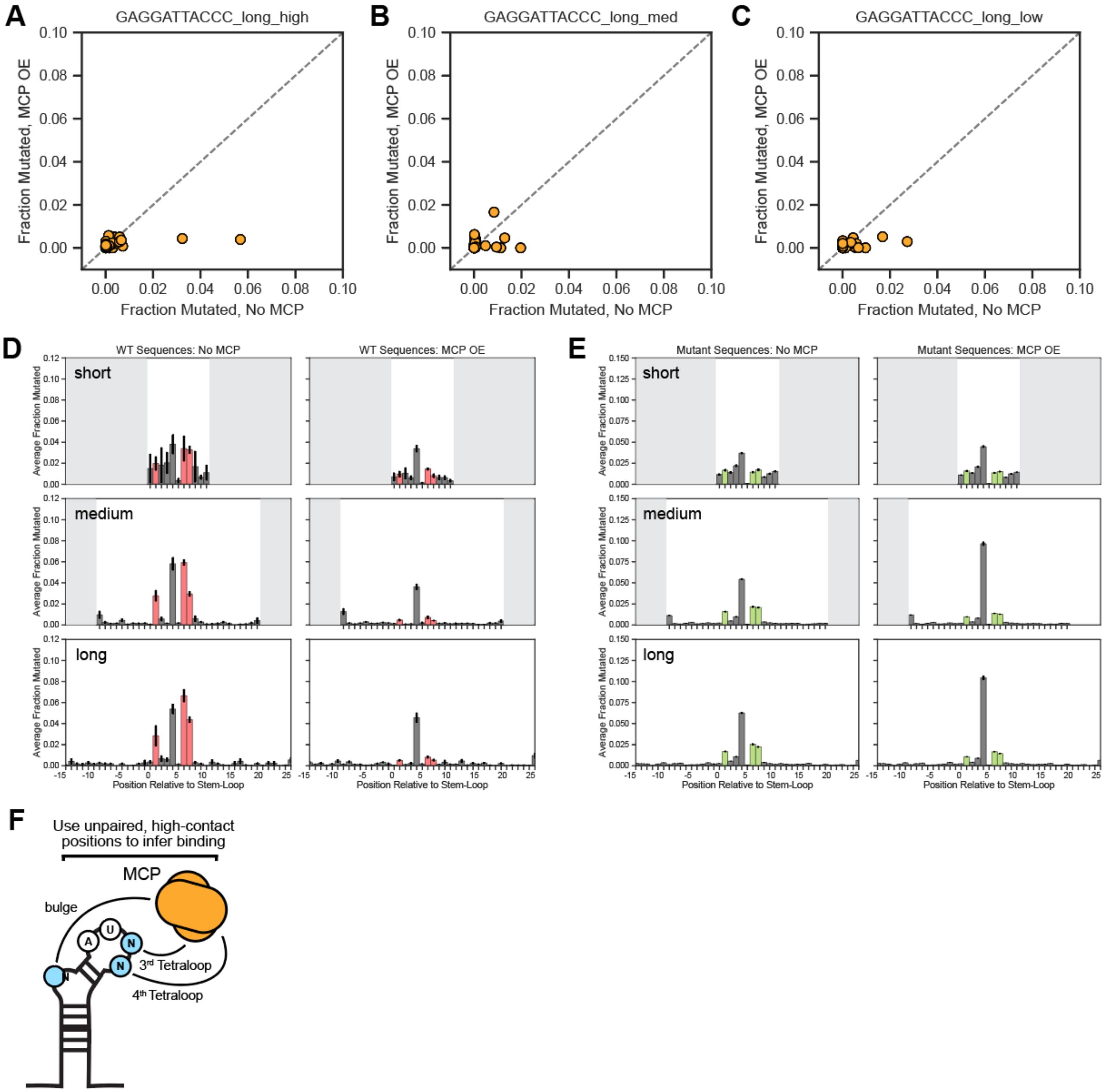


**Figure S3. DMS-MaPseq measurements of library members with changing stem lengths and GC content, related to Figure 2.** (A-C) Comparison of DMS-MaPseq signal at the same positions in the absence (x-axis) and presence (y-axis) of MCP, for the wild-type-like sequence GAGGAUUACCC with a long stem and high (A, 80%), medium (B, 50%), or low (C, 30%) GC content. (D) Accessibility at all positions of all wild-type MS2 library sequences with short (top row), medium (middle row), or long (bottom row) stems, in the absence (left) and presence (right) of MCP, as measured by the DMS-MaPseq mutation fraction at each position. (E) Accessibility at all positions of all mutant MS2 library sequences with short (top row), medium (middle row), or long (bottom row) stems, in the absence (left) and presence (right) of MCP, as measured by the DMS-MaPseq mutation fraction at each position. (F) Schematic of binding metric calculation. Normalized difference values at the three positions that directly contact MCP - the bulge, third tetraloop, and fourth tetraloop positions - are averaged for an “average normalized difference” and qualitative metric to compare binding strengths across library members.

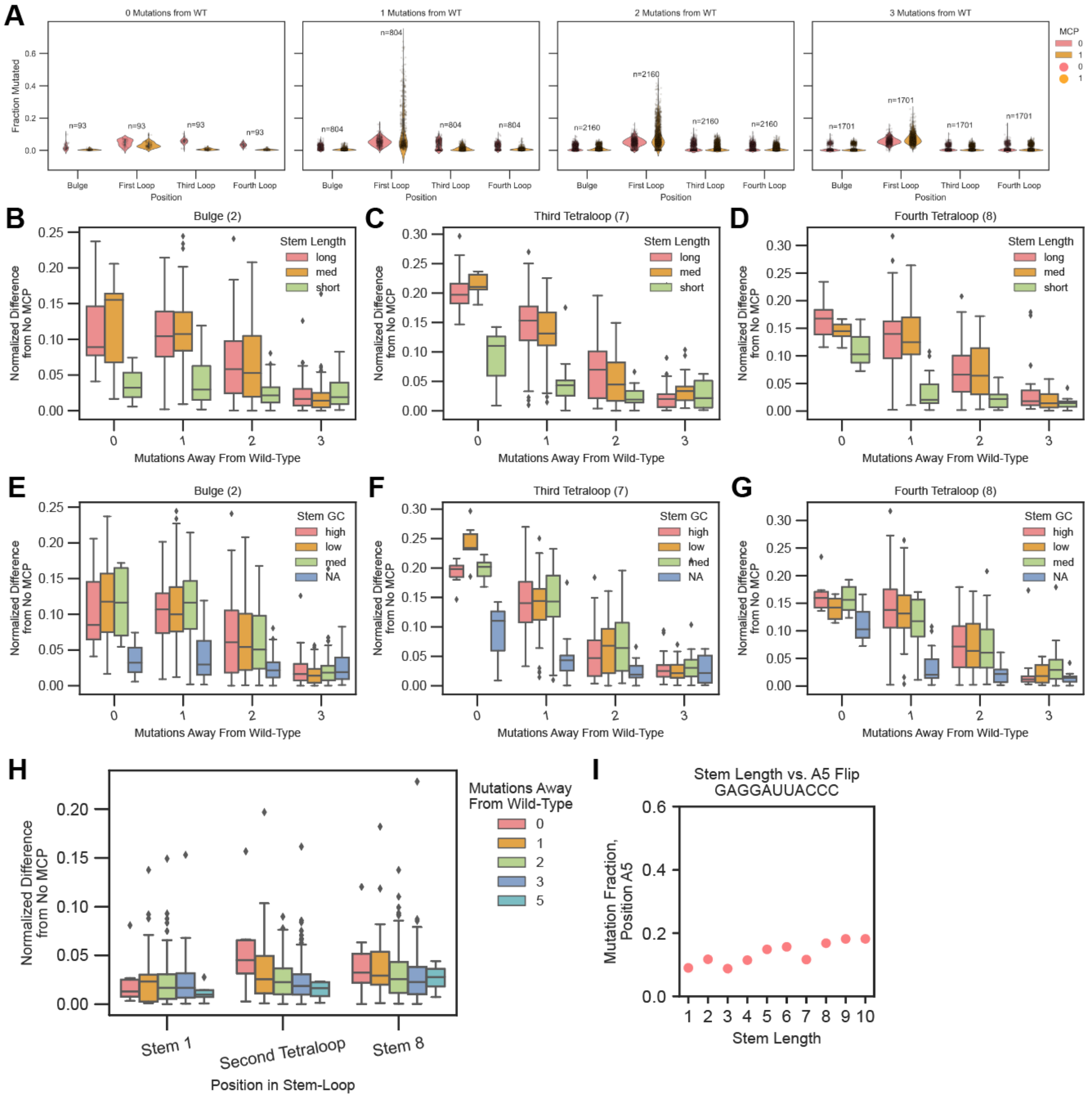


**Figure S4. Mutational distance decreases binding affinity, related to Figure 3.** (A) Fraction mutated values for samples without (pink) and with (orange) MCP, at the bulge position and the first, third, and fourth loop positions in the MS2 tetraloop. L-R, samples with 0, 1, 2, and 3 mutations away from the wild-type consensus. (B-D) Normalized difference in DMS-MaP signal at the bulge (B), third tetraloop (C), and fourth tetraloop (D) positions, separated by the number of mutations that differ from wild-type MS2, and colored by stem length (long: 15bp, medium: 9bp, short: 0bp). Boxes show first to third quartiles, whiskers show 1.5 x IQR, and outliers are shown as diamonds. (E-G) Normalized difference in DMS-MaP signal at the bulge (E), third tetraloop (F), and fourth tetraloop (G) positions, separated by the number of mutations that differ from wild-type MS2, and colored by stem GC content (high: 80%, medium: 50%, low: 30%, NA: 0bp stem). (H) Normalized difference in DMS-MaPseq signal at control positions: the first position in the stem, the second tetraloop position that contacts MCP less consistently, and the eighth stem position 3' to the loop. Sequences are separated and colored by the number of mutations in a sequence that differ from the wild-type MS2. (I) DMS-MaPseq measurements of first tetraloop position A5 for a set of wild-type loop sequences (GAGGAUuACCC) with successively longer stems. Y-axis, DMS-MaPseq mutation fraction at position A5; x-axis, stem length of stem-loop sequence.

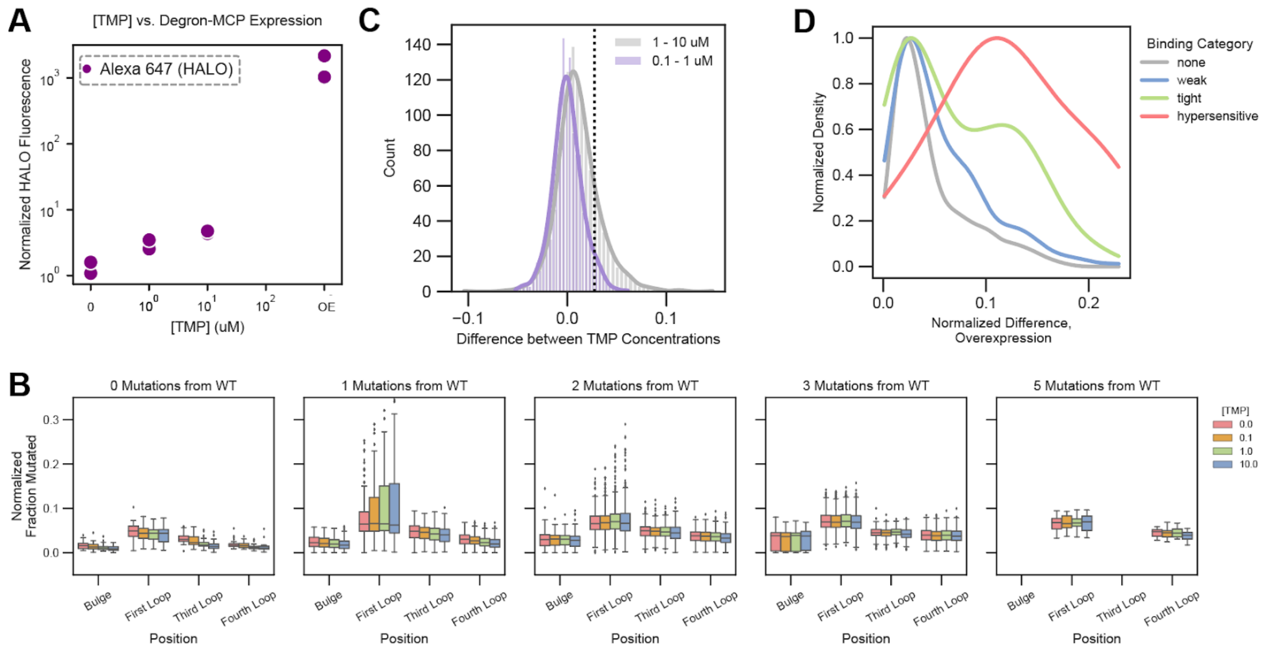


**Figure S5. Details of DHFR-MCP fusion and TMP dosing with DMS-MaPseq experiments, related to Figure 4.** (A) Concentration of DHFR-MCP fusion at different TMP doses, as measured by JaneliaTag-Alexa647 staining of the HALO ligand directly fused to MCP. X-axis, TMP concentration in µM; OE = overexpression of a HaloTag-MCP construct with no DHFR. Y-axis, HaloTag fluorescence normalized to a stained but non-expressing control. (B) Fraction mutated values normalized within each replicate for samples at the bulge position and the first, third, and fourth loop positions in the MS2 tetraloop at different TMP doses. Sequences are separated and colored by TMP dose (0, 0.1, 1, and 10 µM). L-R, samples with 0, 1, 2, 3, and 5 mutations away from the wild-type consensus. Boxes show first to third quartiles, whiskers show 1.5 x IQR, and outliers are shown as diamonds. (C) Distribution of difference values at different TMP doses for average binding values of each sequence in the MS2 library. The averaged normalized difference to 0 TMP (across the bulge, third tetraloop, and fourth tetraloop positions) is taken for each sequence at 3 TMP doses, then the differences calculated between 0.1 and 1 µM TMP (purple) and 1 and 10 µM (purple). Vertical line shows the 95th percentile of the 0.1-1 µM distribution (purple) and the significance cutoff. (D) Distribution of average normalized difference (binding) values in the MCP overexpression experiments shown in Figure 1-3, colored by determined binding category using TMP dosing information as in Figure 4F-4G.

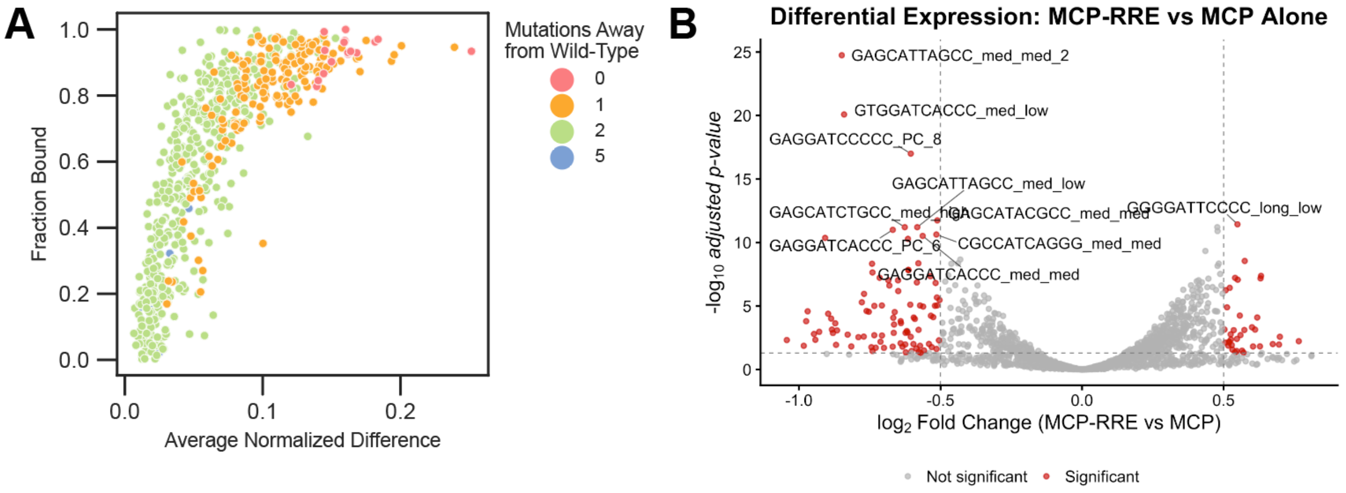


**Figure S6. Transformation of DMS-MaPseq values to fraction occupancy predicts RNA degradation, related to Figure 5.** (A) Comparison of average normalized difference values (x-axis) and transformed fraction bound values (y-axis) for all measurable sequences in the MS2 library (medium and long stem only, removing un-like bases). Sequences are colored by the number of mutations in a sequence that differ from the wild-type MS2. (B) Differential expression of all MS2 library members as measured by RNA-seq during overexpression with MCP alone or the degrading synthetic MCP-RRE fusion. X-axis, relative RNA abundance in $\log_2$FoldChange; smaller values mean lower abundance in the MCP-RRE overexpression sample. Y-axis, $-\log_{10}$ adjusted p-value for each sequence. Significantly differentially expressed transcripts, with a cut off of +/- 0.5 $\log_2$FoldChange and $-\log_{10}$ adjusted p-value of 1, are colored in red.

**Table S1. Sample Conditions, related to Supplementary Dataset**

| Sample | MCP Condition | TMP conc. | % DMS (v/v) |
|---|---|---|---|
| **ART295** | MCP-DHFR | 0 uM | 2% |
| **ART296** | MCP-DHFR | 0.1 uM | 2% |
| **ART297** | MCP-DHFR | 1 uM | 2% |
| **ART298** | MCP-DHFR | 10 uM | 2% |
| **ART299** | MCP-DHFR | 0 uM | 2% |
| **ART300** | MCP-DHFR | 0.1 uM | 2% |
| **ART301** | MCP-DHFR | 1 uM | 2% |
| **ART302** | MCP-DHFR | 10 uM | 2% |
| **L304** | n/a | n/a | 2% |
| **L305** | MCP | n/a | 2% |
| **L306** | MCP | n/a | 2% |
| **L311** | MCP | n/a | 0% |
| **L312** | MCP | n/a | 0% |
| **L313** | MCP-RRE | n/a | 0% |
| **L314** | MCP-RRE | n/a | 0% |

**Supplementary Note: High-Throughput RNA Library Design**

We systematically designed the MS2-MCP library to probe how RNA sequence and structure influence RNA-RNA and RNA-protein interactions. The library samples a defined parameter space, enabling controlled, interpretable perturbations and parallel analysis of thousands of RNA variants under uniform experimental conditions.

To maximize accuracy, reproducibility, and interpretability, the library incorporates several design features: (1) **Accurate sequence identification** - Each RNA variant is linked to a unique 12-nt barcode with a minimum Hamming distance of three, enabling reliable discrimination among closely related sequences and minimizing misassignment from sequencing errors.

(2) **Minimized PCR bias** - All constructs share standardized flanking regions and primer-binding sites and are amplified using low-cycle PCR, ensuring read counts reflect biological signal rather than amplification efficiency.

(3) **High signal-to-noise** - Libraries are sequenced at high depth (typically ≥10,000 reads per variant), with measurement uncertainty estimated by bootstrapping sequencing reads.

(4) **Built-in replicability** – RNAs sequences are represented by multiple independent barcodes, providing internal technical replication, and biological replicates across independent experiments capture variability in cellular context.

### METHODS

#### Data and Code Availability

Processed sequence data is available in the **Supplementary Dataset**. The MCP overexpression, DHFR-MCP, and MCP-RRE plasmids are available on Addgene at IDs #242611, #242615, and #242616, respectively. All other requests for data and reagents will be fulfilled by the Lead Contact upon request.

#### Cell lines and cell culture

All experiments were carried out in HEK293T-LentiX (Takara Bio, 632180, female), which were cultured in a controlled humidified incubator at 37°C and 5% CO2 in DMEM (Gibco, 10569069) media supplemented with 10% FBS (Omega Scientific, 20014T) and 1% Penicillin-Streptomycin-Glutamine (Gibco, 10378016). This cell line was not authenticated. All cell lines tested negative for mycoplasma.

#### Generation of stable DHFR-MCP-expressing cells

Lentiviral production was performed by seeding HEK293T Lenti-X cells at 5 x $10^5$ cells per well in 2 mL of DMEM in 6 wells of a 6-well plate. After 24 hours, cells were transfected with 750 ng of an equimolar mixture of three third-generation production plasmids (pMD2.G: Addgene #12259; pRSV-Rev: Addgene #12253; pMDLg/pRRE: Addgene #12251; all gifts from D. Trono) and 750 ng of pAT124 (DHFR-MCP overexpression plasmid). The 4 plasmids were incubated for 15 minutes with 5 µL of polyethylenimine (PEI, Polysciences #23966) before transfection. After 72 hours of incubation, lentivirus was harvested and collected, and supernatant was filtered through 0.45 µM PES filters (CELLTREAT #229749). 12 mL of virus was concentrated into 1 mL using Pierce PES Protein Concentrators (Thermo Fisher #88533), which was added directly to a 10-cm plate of HEK293T cells at 50% confluence. After 48 hours of culture, antibiotic selection was initiated with blasticidin (10 µg/mL, Gibco #A1113903) and infection and selection efficiency were monitored daily with flow cytometry on a Bio-Rad ZE5 Cell Analyzer (Bio-Rad #12004278).

#### MS2 stem-loop library design

Three base wild-type MS2 stem-loops were used for successive mutation: **GAGGAUCACCC**, **CACCAUCAGGG**, and **GAGCAUCAGCC**. All possible single, double, and triple substitutions of the four nucleotides (A,G,C,U) were created for each stem-loop at positions A2 (bulge), C7 (third tetraloop), and A8 (fourth tetraloop) for 192 total stem-loops. Several presumed non-binding sequences were also created by taking the base stem-loops **GAGGAUUACCC**, **CACCAUUAGGG**, and **GAGCAUUAGCC** and substituting the conserved position A5 (first tetraloop) for C, G, and U. Two sequences with the bottom C-G stem of the stem-loop disrupted, **AGAGAUUACCU** and **GGAAAUUAUCC**, were also added as non-binding controls for a total of 206 base stem-loops.

6 stem variants were designed: 9- and 15-bp long with low (30%), medium (50%) or high (80%) GC content. Each loop was added to each stem variant, plus repeated once with no additional stem past the 3 G-C pairs in the loop, for a total of 1,442 members. Finally, a set of 16 stem-loops (the consensus **GAGGAUCACCC** plus all variants at the 7th and 8th positions) added to a progressively longer stem of 1-10 base pairs for a total of 160 additional sequences were created for a total of 1,602 library members.

The final library sequence were constructed as follows: a forward primer-binding sequence for amplification (**TTAAACCGGCCAACATACC**), a C/T shuffled buffer sequence, the forward stem sequence, the loop variant, the reverse stem sequence, another C/T shuffled buffer sequence, a unique 15 nucleotide barcode sequence (no closer than Hamming distance 3 to any other barcode), and a reverse primer-binding sequence **CGCTACTCGTTCCTTTCGA**. The C/T buffer sequences on each side of the stem-loop were variable in length so that each library member was exactly 170 nucleotides long. The library was ordered as a pool from Twist Biosciences.

**Pooled library cloning**

The Twist oligo pool was resuspended to 10 ng/μL in water and the library was PCR amplified using primers specific to the appended primer-binding sequences, above. The reactions were prepared in a pre-PCR hood to reduce contamination. PCR was performed using 10 μL Q5 polymerase buffer (NEB #B9027S), 10 μL High GC Enhancer (NEB #B9028A), 1 μL dNTP mix (NEB #N0447S), 0.5 μL Q5 polymerase (NEB #M0491S), 2.5 μL each of forward and reverse primer (10 μM), and 1 μL of the library pool (10 ng). The following protocol was used: initial denaturation at 98°C for 30s; 6 cycles of 98°C for 10s, 63°C for 30s, and 72°C for 60s; final extension at 72°C for 2 minutes. The amplified library were purified with 0.9X SPRISelect (Beckman Coulter #B23317) and elution in 20 μL.

pcDNA3.1-mCherry (Addge #128744) was double-digested with 10,000 U/mL NotI-HF (NEB #R3189L) and XbaI (NEB #R0145L) for 30 minutes at 37°C, using 1 μL of each enzyme per 5 μg plasmid. After heat inactivation at 65°C for 20 minutes, the digested plasmid was run on a 0.5% TAE gel until a linearized band could be extracted using the QIAquick Gel Extraction Kit (Qiagen #28704). The amplified library was cloned into the digested pcDNA3.1 using Gibson assembly with the NEBuilder HiFi DNA Assembly kit (NEB #E5520S) as follows: 8 Gibson reactions were prepared using 10 μL HiFi kit, 15 ng of PCR product, 100 ng of vector, and nuclease-free water to 20 μL. Reactions were incubated at 50°C for 1 hour, then pooled together and concentrated to 6 μL using the Zymo Clean&Concentrate DNA kit (Zymo #D4004).

2.5 μL of the pooled Gibson assemblies were electroporated into 50 μL of of Endura DUO electrocompetent cells (Lucigen #60242-2) using Gene Pulse Electroporation Cuvettes with a 0.1cm band gap (Bio-Rad #1652089) and a Gene Pulser Xcell Total System (Bio-Rad #1652660) under the following conditions: 1.8kV, 10 uF, 600 Ω, and 0.1 cm distance. Cells were recovered in 2 mL of 37°C SOC recovery medium (NEB #B9020S) at 37°C for 1 hour, after which they were added directly to 500 mL LB with 100 μg/mL carbenicillin. Cells were incubated overnight at 30°C, after which the plasmid pool was extracted using the Qiagen Plasmid Maxi Kit (Qiagen #12162) and library quality was assessed using Illumina sequencing after PCR amplification from the plasmid pool.

**Transfection of stem-loop library and MCP constructs**

Libraries and overexpression plasmids were transfected into wild-type or DHFR-MCP-expressing HEK293T cells using Lipofectamine 3000 following the manufacturer's protocol. Briefly, one 10-cm plate for each biological replicate and experimental condition was seeded with 4 x $10^6$ HEK293T cells. 24 hours later, 7 μg of library alone or 7 μg of library plus 7 μg of pAT031 (MCP overexpression plasmid) or pAT121 (MCP-RRE degrader plasmid) were transfected onto each plate using 21.7 μL of Lipofectamine 3000 and 28 μL of P3000 reagent. After 48 hours, cells were processed either for DMS-MaPseq or RNA-seq (see below). For experiments with stable DHFR-MCP-expressing HEK293T cells, TMP was added to a given concentration (0.1, 1, or 10 μM final) 24 hours before library transfection to allow for MCP expression, then refreshed every 24 hours to with media changes to maintain constant drug concentration through the course of the experiment.

**DMS-MaPseq for detection of RNA structure and MCP binding**

10 mL of DMEM per 10-cm plate of HEK293T cells was warmed in preparation for DMS modification. 10 mL per sample of each of 30% 2-mercaptoethanol (Sigma-Aldrich #M6250) in PBS (Thermo Fisher #14190250), 10% 2-mercaptoethanol in PBS, and PBS alone were chilled on ice in a chemical fume hood. Cells in 10-cm plates were brought into a tissue culture hood and media was aspirated. 2% v/v of DMS (dimethyl sulfate, Thermo Fisher #430831000) was added to 10 mL DMEM per sample using a needle or under hypoxic conditions, quickly vortexed, and immediately added to cells. Cells were returned to a 37°C incubator for exactly 3 minutes, after which they were brought to the chemical fume hood and quenched with 10 mL per sample of ice-cold 30% 2-mercaptoethanol in PBS. After quenching, cells were moved to a 50 mL conical tube and centrifuged for 5 minutes at 1000xg at 4°C. Pellets were resuspended in ice-cold 10% 2-mercaptoethanol in PBS and re-centrifuged for 5 minutes at 1000xg at 4°C. Finally, pellets were resuspended in ice-cold PBS and centrifuged for 5 minutes at

1000xg at 4°C, after which they were resuspended in 1 mL Trizol (Thermo Fisher #15596026) and frozen at -80°C.

**Targeted RNA-seq for detection of stem-loop library member abundances**

The stem-loop library and either pAT031 (MCP overexpression plasmid) or pAT121 (MCP-RRE overexpression plasmid) were expressed in cells for 48 hours, after which each 10-cm plate was harvested by centrifugation. Library prep was performed as follows with DMS samples, below.

**RNA library preparation for Illumina sequencing for DMS-MapSeq and RNA-seq samples**

RNA was extracted using Trizol-chloroform extraction as follows: 200 µL chloroform (VWR #J67241-AP) was added to each Trizol sample, vortexed well, and centrifuged for 15 minutes at 15,000xg. The upper (aqueous) layer was mixed with an equal volume of 100% ethanol, then added to an RNEasy column from the RNEasy Mini kit (Qiagen #74106). RNA was extracted with the kit by following the rest of the protocol per manufacturer's instructions. 5 µg of each RNA sample was depleted of rRNA species by incubation with 5 µg subtraction oligos in a total volume of 1x hybridization buffer (200 mM NaCl, 100 mM Tris pH 7.5) for 1 minute at 68°C, then the temperature ramped down at a rate of 1°C/min to 50°C. 10 µL RNase H buffer and 1 µL thermostable RNase H (NEB #M0523S) were added directly to each reaction along with 59 µL nuclease-free water. Reactions were incubated for 20 minutes at 50°C, then purified using the Zymo RNA Clean&Concentrate-5 kit (Zymo #R1013) with pre-column DNase I treatment as included in the kit.

1 µg of each rRNA-depleted RNA sample was then diluted to 11 µL in nuclease-free water and incubated with 1 µL of 10 µM reverse primer, designed to bind the reverse primer-binding sites on the library members, and 1 µL dNTP solution mix (10 mM, NEB #N0447S) for 5 minutes at 65°C. 4 µL Induro RT reaction buffer (NEB #B0681), 0.5 µL RNase inhibitor (NEB #M0314), 1 µL Induro Reverse Transcriptase (NEB #M0681S), and 1.5 µL nuclease-free water were added to each reaction, which were then incubated for 90 minutes at 55°C. Following incubation, 1 µL of 4 mM NaOH was added to each reaction and incubated for 3 minutes at 95°C to quench reverse transcription. The resultant single-stranded DNA samples were purified using the Zymo DNA Clean&Concentrate-5 kit (Zymo #D4004) following the protocol for ssDNA.

Samples were amplified with PCR with the following protocol: 10 µL high GC enhancer, 0.5 µL Q5 polymerase, 1 µL dNTP mix, 10 µL Q5 buffer, 2.5 µL each of 10 µM forward and reverse primers that bind the sites designed on the library members, 3 µL each cDNA product, and 20.5 µL nuclease-free water. They were amplified as follows: initial denaturation at 98°C for 30s; 25 cycles of 98°C for 10s, 63°C for 30s, and 72°C for 60s; final extension at 72°C for 2 minutes. Samples were purified with the Zymo DNA Clean&Concentrate-5 kit and eluted in 10 µL nuclease-free water. 250 ng of purified dsDNA product was prepared for Illumina sequencing following the NEBNext Ultra II DNA Library Prep Kit for Illumina (NEB #E7645L) following manufacturer's instructions. Briefly, 250 ng of DNA was diluted to 50 µL and incubated with DNA End Prep Enzyme Mix and Buffer (#E76746) for 30 minutes at 20°C followed by 30 minutes at 65°C, then added to 2.5 µL undiluted Illumina adaptor, Ligation Master Mix (#E7648), and Ligation Enhancer (#E7374) for 15 minutes at 20°C. USER enzyme was used to digest closed-loop adapters for 15 minutes at 37°C, and reactions were purified with a double-sided 0.5X followed by 0.25X SPRISelect purification. Finally, samples were amplified with Q5 Ultra II Master Mix (#E7649) using NEBNext Multiplex Oligos for Illumina (NEB #E7730) before final purification with 0.9X SPRISelect cleanup. Libraries were quantified using the Qubit dsDNA HS Assay Kit (Thermo #Q33231) on a Qubit 4 Fluorometer (Fisher #Q33238) and assessed for purity on an Agilent TapeStation (Agilent #G2964AA) before being sequenced on an Illumina NovaSeq with 2x150 cycles.

**DMS-MaPseq data analysis**

The FASTQ files were processed using SEISMIC-RNA (version 0.24.2)[3] as previously described.[4] Briefly, the function *wf* (for workflow) was executed including *-x* for 2 separate FASTQ files of paired-end reads and a reference FASTA file containing the sequence of each reference in the library. Processed data was further analyzed and visualized using custom Python scripts. The data was replotted in GraphPad Prism 10 to increase the resolution for publication.

**RNA-seq data analysis**

Processing scripts were modified from https://github.com/bintulab/Viral_Ludwig_2022/tree/main/final_April-2023/RNA-seq%20Scripts. Briefly, reads were demultiplexed with bcl2fastq and aligned to a reference transcriptome with hisat2. The reference transcriptome was built from a .fasta file containing the sequences of all MS2 stem-loop library members using hisat2-build. Output SAM files were converted to BAM files, sorted, and indexed using samtools. Differential expression analysis was performed in R with the Bioconductor DESeq2 package[80] and a set of custom R scripts, based on the following tutorial: http://bioconductor.org/help/course-materials/2016/CSAMA/lab-3-rnaseq/rnaseq_gene_CSAMA2016.pdf. Further analyses were performed and visualized using custom Python scripts.

**Derivation of fraction bound from DMS-MaPseq mutation data**

The mutation fraction at each position *j* as measured by DMS-MaPseq for a given RNA sequence can be represented by the initial rate equation. The DMS modification of the unfolded and folded RNA states (while unbound) follows second-order rate kinetics, in which the reaction is dependent both on the number of RNA molecules existing in that state and on the concentration of DMS. In the bound RNA state, the modification rate is considered negligible due to shielding by MCP, which reduces the pseudo-second-order rate constant to near zero (pseudo-second order reaction, $k_{DMS,\,B,\,j} = 0$).

The observed mutation fraction in each experiment for a given position is thus:

$$MF_{j,observed} = \left(P_{unfolded}\, k_{DMS,U,j}[DMS] + P_{folded}\, k_{DMS,F,j}[DMS] + P_{bound}\, k_{DMS,B,j}[DMS]\right)t$$

Using the stem length and folding analyses in **Figure 2**, we have determined that every library member with a medium or long stem is folded in cells, evidenced by a lack of DMS reactivity at the paired stem bases. By restricting our analyses to only those library members, we know that the fraction of unfolded RNAs is negligible ($P_{unfolded} \approx 0$). We also use only RNA positions that directly contact MCP to approximate binding, for which the rate of DMS reactivity ($k_{\mathrm{DMS,B}}$) is essentially zero when bound to MCP. We can then simplify:

$$MF_{j,\,observed} = k_{DMS,\,F,\,j}[DMS]t$$

We also assume that all measured positions in the absence of MCP can be represented by $P_{folded} = 1$ (since we selected library members that fold completely), which means that

$$MF_{j,\,observed}^{-MCP} = k_{DMS,\,F,\,j}[DMS]t$$

This indicates that each mutation fraction that we measure at a binding position is directly related to the probability of that sequence existing in a folded, but not bound, state. We solve for $P_{folded}$:

$$P_{folded} = \frac{MF_{j,\,observed}}{k_{DMS,F,j}\,[DMS]t}$$

When the sequence is completely folded and bound by MCP, $P_{folded} = 0$; inversely, $P_{folded} = 1$ indicates a complete lack of protein binding at that position. We assume that the wild-type sequence **GAGGAUCACCC_long_med**, shown in **Figure 1,** is completely bound and assign it as a reference wild-type sequence. We can then calculate a change in abundance between the folded and unbound fraction for a mutant RNA compared to the WT, $\Delta P_{folded}$:

$$\Delta P_{folded} = P_{folded}^{WT} - P_{folded}^{mut} = \frac{MF_{j,\,observed}^{WT}}{k_{DMS,F,j}\,[DMS]t} - \frac{MF_{j,\,observed}^{mut}}{k_{DMS,F,j}\,[DMS]t}$$

We remove all sequences with $\Delta P_{folded} > 1$, as these are not physically possible values and represent measurement noise. Importantly, we also reduce our analysis to only common bases shared between the reference wild-type and each mutant sequence to ensure a constant $k_{DMS}$, which is variable per base, although this removes all sequences in the library that are mutated at all of position 2, 7, and 8.

This leaves us with a final simplification of

$$\Delta P_{folded} = \frac{MF_{j,observed}^{WT,+MCP}}{MF_{j,observed}^{WT,-MCP}} - \frac{MF_{j,observed}^{mut,+MCP}}{MF_{j,observed}^{mut,-MCP}}$$

Since we assume these RNAs are entirely folded, the fraction bound is simply:

$$f_{bound} = \Delta P_{bound} = 1 - \Delta P_{folded}$$
